# Dopamine-driven Increase in IL-1β in Myeloid Cells is Mediated by Differential Dopamine Receptor Expression and Exacerbated by HIV

**DOI:** 10.1101/2024.06.09.598137

**Authors:** Stephanie M. Matt, Rachel Nolan, Samyuktha Manikandan, Yash Agarwal, Breana Channer, Oluwatofunmi Oteju, Marzieh Daniali, Joanna A. Canagarajah, Teresa LuPone, Krisna Mompho, Kaitlyn Runner, Emily Nickoloff-Bybel, Benjamin Li, Meng Niu, Johannes C. M. Schlachetzki, Howard S. Fox, Peter J. Gaskill

## Abstract

The catecholamine neurotransmitter dopamine is classically known for regulation of central nervous system (CNS) functions such as reward, movement, and cognition. Increasing evidence also indicates that dopamine regulates critical functions in peripheral organs and is an important immunoregulatory factor. We have previously shown that dopamine increases NF-κB activity, inflammasome activation, and the production of inflammatory cytokines such as IL-1β in human macrophages. As myeloid lineage cells are central to the initiation and resolution of acute inflammatory responses, dopamine-mediated dysregulation of these functions could both impair the innate immune response and exacerbate chronic inflammation. However, the exact pathways by which dopamine drives myeloid inflammation are not well defined, and studies in both rodent and human systems indicate that dopamine can impact the production of inflammatory mediators through both D1-like dopamine receptors (DRD1, DRD5) and D2-like dopamine receptors (DRD2, DRD3, and DRD4). Therefore, we hypothesized that dopamine-mediated production of IL-1β in myeloid cells is regulated by the ratio of different dopamine receptors that are activated. Our data in primary human monocyte-derived macrophages (hMDM) indicate that DRD1 expression is necessary for dopamine-mediated increases in IL-1β, and that changes in the expression of DRD2 and other dopamine receptors can alter the magnitude of the dopamine-mediated increase in IL-1β. Mature hMDM have a high D1-like to D2-like receptor ratio, which is different relative to monocytes and peripheral blood mononuclear cells (PBMCs). We further confirm in human microglia cell lines that a high ratio of D1-like to D2-like receptors promotes dopamine-induced increases in IL-1β gene and protein expression using pharmacological inhibition or overexpression of dopamine receptors. RNA-sequencing of dopamine-treated microglia shows that genes encoding functions in IL-1β signaling pathways, microglia activation, and neurotransmission increased with dopamine treatment. Finally, using HIV as an example of a chronic inflammatory disease that is substantively worsened by comorbid substance use disorders (SUDs) that impact dopaminergic signaling, we show increased effects of dopamine on inflammasome activation and IL-1β in the presence of HIV in both human macrophages and microglia. These data suggest that use of addictive substances and dopamine-modulating therapeutics could dysregulate the innate inflammatory response and exacerbate chronic neuroimmunological conditions like HIV. Thus, a detailed understanding of dopamine-mediated changes in inflammation, in particular pathways regulating IL-1β, will be critical to effectively tailor medication regimens.

## 1. Introduction

Dopamine is an endogenous catecholamine neurotransmitter that has a well-established role in regulating motor control, cognition, motivation, and reward in the central nervous system (CNS). Careful regulation of dopamine concentrations and bioavailability is crucial for normal CNS function and protection against the development of neurological diseases, such as Parkinson’s disease (hypodopaminergic) or schizophrenia (hyperdopaminergic). However, dopamine is not only found in the CNS, but also has a unique distribution pattern throughout the circulatory system and in many peripheral tissues, where it is important to both neuronal and nonneuronal processes (Matt and Gaskill 2019, Channer, Matt et al. 2022). For example, changes in peripheral dopamine can regulate metabolism, gastrointestinal motility, kidney function, blood pressure maintenance, and many other functions (Eisenhofer, Aneman et al. 1997, Fitzgerald and Dinan 2008, Armando, Villar et al. 2011, Harris and Zhang 2012, Blum, Thanos et al. 2014). In addition, there is increasing recognition that dopamine has a substantial regulatory role in immune cell function in both the CNS and periphery.

Dopaminergic immunomodulation affects innate immunity by modulating the function of myeloid lineage cells such as microglia and macrophages (Channer, Matt et al. 2022), which are the most common immune cells in the CNS (Ransohoff and Cardona 2010, Herz, Filiano et al. 2017, Prinz, Masuda et al. 2021). Microglia, which are exposed to high levels of dopamine in specific brain regions, exhibit changes in neuroinflammatory cytokine production, phagocytosis, and formation of extracellular traps (Huck, Freyer et al. 2015, Dominguez-Meijide, Rodriguez-Perez et al. 2017, Kopec, Smith et al. 2017, Fan, Chen et al. 2018, Agrawal, Sharma et al. 2021). In the periphery, circulating monocytes and differentiated macrophages within tissues that fulfill niche-specific functions can also respond to dopamine. Data show primary human and rodent monocytes/macrophages can take up, store, and produce dopamine (Brown, Meyers et al. 2003, Flierl, Rittirsch et al. 2009, Kokkinou, Fragoulis et al. 2009, Gopinath, Badov et al. 2021, Mackie, Gopinath et al. 2022), and dopamine can modulate myeloid inflammatory signaling. We have previously published that dopamine increases the production of a number of inflammatory cytokines and chemokines in primary human macrophages (Nolan, Muir et al. 2018). These effects, at least in part, are induced by dopamine receptor-mediated activation of NF-κB, resulting in the induction of its downstream targets including the NLRP3 inflammasome and the inflammatory IL-1 cytokine family including IL-1β and IL-18 (Nolan, Reeb et al. 2020). These data are supported by additional studies showing dopamine receptor activity is inflammatory (Nakano, Higashi et al. 2008, Nakano, Yamaoka et al. 2011, Fan, Chen et al. 2018), although other studies have also shown dopamine to have anti-inflammatory effects on NF-κB and NLRP3 (Yan, Jiang et al. 2015, Wang, Nowrangi et al. 2018, Liu and Ding 2019, Han, Ni et al. 2020, Yue, Wang et al. 2021).

Fluctuations in extracellular dopamine concentrations could occur in response to the use of dopamine altering therapeutics, or during the use of addictive substances such as stimulants or opioids (Brannan, Martinez-Tica et al. 1993, Ichikawa and Meltzer 1995, Seeman and Madras 2002, Li, Ichikawa et al. 2003, Li, Ichikawa et al. 2004, Ashok, Mizuno et al. 2017). Different classes of substances of misuse act via distinct mechanisms, but all addictive substances increase CNS dopamine *in vivo,* and dopamine signaling plays a significant role in substance use disorders (SUDs) (Di Chiara 1999, Di Chiara 2000, Pierce and Kumaresan 2006, Bloomfield, Ashok et al. 2016, Kokkinou, Ashok et al. 2018). In both humans and animal models, psychostimulants such as methamphetamine and cocaine promote myeloid neuroinflammation *in vivo* and *in vitro* (Cui, Shurtleff et al. 2014, Frank, Adhikary et al. 2016, Lacagnina, Rivera et al. 2017, Kohno, Link et al. 2019, Namba, Leyrer-Jackson et al. 2021). Stimulant use increases the total area of CNS exposed to aberrant dopamine levels (Cragg, Nicholson et al. 2001, Robinson, Venton et al. 2003, Venton, Zhang et al. 2003, Rice and Cragg 2008, Sulzer, Cragg et al. 2016), thereby increasing the number of myeloid cells exposed to those elevated dopamine concentrations. This could increase inflammatory cytokine production in myeloid cells in dopamine-rich regions such as the nucleus accumbens and ventral tegmental area, explaining the inflammatory effects in these regions after drug exposure (Cearley, Blindheim et al. 2011, Northcutt, Hutchinson et al. 2015, Frank, Adhikary et al. 2016, Brown, Levis et al. 2018). Critically, these effects are not observed *ex vivo* in isolated microglia (Cearley, Blindheim et al. 2011, Frank, Adhikary et al. 2016), suggesting that the neuronal release of dopamine in response to these stimulants may be required to mediate the inflammatory effects in myeloid cells.

Most types of immune cells can respond to dopamine through their expression of dopamine receptors and other dopaminergic proteins, which may lead to changes in immune function (Channer, Matt et al. 2022). Dopamine primarily acts via 5 subtypes of G-protein coupled receptors known as D1-like (DRD1 and DRD5) and D2-like (DRD2, DRD3, and DRD4) dopamine receptors (Beaulieu and Gainetdinov 2011). Nonetheless, the varying affinities of these receptors for dopamine (Gaskill, Calderon et al. 2013) suggests that the ratio of dopamine receptors is important. The balance of D1-like vs. D2-like receptor signaling – and the associated outcomes - could change drastically depending on extracellular dopamine concentrations (Pacheco 2017, Martel and Gatti McArthur 2020). Overall, this suggests that dopaminergic immunomodulation could be a common mechanism by which substances of misuse could influence the development of chronic inflammation and risk for neurological disease. In addition, many clinically relevant therapeutics that are dopamine modifying agents, including but not limited to bromocriptine (type 2 diabetes), carbidopa (Parkinson’s disease), bupropion (depression), and aripiprazole (schizophrenia) can also impact inflammation (Sobiś, Rykaczewska-Czerwińska et al. 2015, Jha, Minhajuddin et al. 2017, Zhu, Lemos et al. 2017, Cincotta, Cersosimo et al. 2022), potentially through their effects on the dopamine system in immune cells. However, the specific dopamine receptors that drive these effects in human myeloid cells remain inconsistent and poorly understood.

To address this knowledge gap, we examined the role of dopamine receptors in the regulation of IL-1β in several types of human myeloid cells. In myeloid cells, this cytokine is linked to cell activation and is associated with many human infectious and non-infectious neurological diseases (Stojakovic, Paz-Filho et al. 2017, Liu and Quan 2018, Mendiola and Cardona 2018, Rivera-Escalera, Pinney et al. 2019). IL-1β is a master regulator of inflammation and plays multifaceted roles in the periphery, but the CNS is especially sensitive to IL-1β signaling because multiple CNS cell types express receptors for IL-1β (Walsh, Reinke et al. 2014, Liu, Nemeth et al. 2019, Nemeth and Quan 2021). Due to this sensitivity, the secretion of the active form of IL-1β is tightly regulated by inflammasomes, which are multi-protein complexes that control the maturation and release of IL-1 family cytokines (Mariathasan and Monack 2007, Chan and Schroder 2020). In the context of dopamine and dopamine-inducing drugs, IL-1β can be directly increased by substances of misuse through interactions with toll-like receptor 4 (TLR4) (Northcutt, Hutchinson et al. 2015). IL-1β can also uniquely alter the function and expression of dopamine receptors as well as dopamine biosynthesis rate and storage *in vivo* and *in vitro* relative to other cytokines (Shintani, Kanba et al. 1993, Abreu, Llorente et al. 1994, Lacosta, Merali et al. 1998, Ling, Potter et al. 1998), suggesting a greater connection between these two systems than is currently defined. The connections between IL-1β and the dopaminergic system have primarily been studied in whole brain regions and neuronal populations, leaving the relationship between IL-1β and the myeloid dopaminergic system unclear. We examined dopamine receptor-mediated IL-1β regulation in primary human macrophages (hMDM) and microglia, using human microglial cell lines as well as inducible pluripotent stem cell-derived human microglia. Our data in hMDM indicate that while expression of DRD1 is required for dopamine to increase IL-1β, levels of D2-like receptors can modulate the magnitude of this effect. These findings are supported by studies in microglial cell lines confirming the importance of both types of receptors to IL-1β production in myeloid cells. RNA sequencing further corroborated our results and showed that dopamine mediates changes in pathways related to IL-1β signaling in human microglia.

We then examined HIV-infected cells as an example of a neuroimmunological disease that could be exacerbated by SUDs, which are a common comorbidity in people living with HIV (PLWH) (Durvasula and Miller 2014, Saylor, Dickens et al. 2016, Chilunda, Calderon et al. 2019). HIV enters the CNS within weeks of initial infection, infecting long-lived myeloid cells such as microglia and CNS-resident macrophages, which are the primary targets for HIV in this compartment (Avalos, Price et al. 2016, Clayton, Garcia et al. 2017, Wallet, De Rovere et al. 2019). These cells are also major regulators of the innate immune response, driving a chronic, neuroinflammatory state that leads to a variety of neuropathologic effects known as neuroHIV (Borrajo, Spuch et al. 2021, Borrajo López, Penedo et al. 2021, Muzio, Viotti et al. 2021), which remains prevalent in individuals on antiretroviral therapy (ART) (Saylor, Dickens et al. 2016).

Even with effective ART, infected individuals show substantial pathology in dopamine-rich brain regions, including volume decline, increased myeloid cell numbers and activity, and neuronal dysfunction (Becker, Sanders et al. 2011, Ipser, Brown et al. 2015, Kallianpur, Jahanshad et al. 2020). These data are corroborated by studies in SIV-infected rhesus macaques in which direct increases in CNS dopamine levels via treatment with L-DOPA, methamphetamine, or selegiline significantly enhance neuropathology, increasing viral load, inflammation in dopamine rich brain regions, and the proportion of SIV-infected myeloid cells (Czub, Koutsilieri et al. 2001, Czub, Czub et al. 2004, Marcondes, Flynn et al. 2010, Najera, Bustamante et al. 2016, Niu, Morsey et al. 2020). We have previously shown that SIV infection alone increases dopamine levels in caudate and frontal cortex, and that selegiline-mediated increases in dopamine in SIV-infected macaques decrease the expression of antiviral genes (Emanuel, Runner et al. 2022).

Notably, HIV is a prime example of a pathological condition where the role of IL-1β is well established, as it is elevated in the CNS and serum during HIV and SIV (Zhao, Kim et al. 2001, Walsh, Reinke et al. 2014, Triantafilou, Ward et al. 2021, Farhadian, Lindenbaum et al. 2022, Yaseen, Abuharfeil et al. 2023). NLRP3 inflammasomes have also been shown to be upregulated by HIV in CNS and peripheral myeloid cells (Walsh, Reinke et al. 2014, Zhang, Mosoian et al. 2019, Triantafilou, Ward et al. 2021). Both inflammasome activation and IL-1β release are seen in human and rodent macrophages and mixed glial cultures in response to live virus or soluble HIV proteins such as Tat, gp120, and Vpr (Clouse, Cosentino et al. 1991, Koka, He et al. 1995, Cheung, Ravyn et al. 2008, Walsh, Reinke et al. 2014, Triantafilou, Ward et al. 2021). This indicates that productive viral replication of cells is not a requirement of induction of this inflammation. As we hypothesized, the data indicate that the effects of dopamine on IL-1β and inflammasome components are altered in the presence of HIV infection. These findings suggest an unexplored mechanism underlying dopamine-mediated myeloid cell activation associated with signaling and processing of IL-1β, and provide a potential target for the development of therapeutics aimed at managing inflammation in disease.

## 2. Methods

### 2.1 Reagents

DMEM (cat # 10569044) and RPMI-1640 (cat # 11875119) medium, sodium pyruvate (cat # 11360070), trypsin, and penicillin/streptomycin (P/S) (cat # 15140163) were from Invitrogen (ThermoFisher, Carlsbad, CA, USA). Bovine serum albumin (BSA) (cat # BP1600100), glycine (cat # G48500), 4% Paraformaldehyde solution (cat # AAJ19943K2), blasticidin (cat # R21001), Zeocin (cat # R25001), and calcium phosphate transfection kits (cat # K278001) were from Fisher Scientific (Waltham, MA, USA). Hydroxyethyl piperazineethanesulfonic acid (HEPES) (cat # BP2991), Tween (cat # NC0689073), dimethyl sulfoxide (DMSO) (cat # 472301), Poly I:C (cat # P1530), and dopamine hydrochloride (cat # H8502) were obtained from Sigma-Aldrich (St. Louis, MO, USA). Fetal calf serum (FBS) was from Corning (cat # MT35010CV), human AB serum was from Gemini Bio-Products (cat # 100-512), and PEG-It Virus Precipitation Solution (cat # LV825A-1) was from System Biosciences. Macrophage colony stimulating factor (M-CSF) (cat # 300-25), IL-34 (cat # 200-34), and TGF-β1 (cat # 100-21) were from Peprotech (Rocky Hill, NJ, USA). TaqMan Fast Universal Master Mix, and PCR assay probes for DRD1-5 (Hs00265245_s1, Hs00241436_m1, Hs00364455_m1, Hs00609526_m1, Hs00361234_s1), IL-1β (Hs01555410_m1), NLRP3 (Hs00918082_m1), NLRC4 (Hs00368367_m1), AIM2 (Hs00915710_m1), NLRP1 (Hs00248187_m1), NLRC5 (Hs01072123_m1), IL-18 (Hs01038788_m1), IL1rn (Hs00893626_m1), Pycard (Hs01547324_gH), Casp1 (Hs00354836_m1), and 18s (4319413E) genes were purchased from Applied Biosystems (ThermoFisher, Waltham, MA, USA). Flupentixol dihydrochloride (cat # 4057/50) was from R&D Systems (Minneapolis, MN, USA), and was diluted to a stock concentration of 100mM in dH_2_O and aliquoted and stored at -20°C until use.

Dopamine hydrochloride (DA) was resuspended in dH_2_O at a stock concentration of 10 mM that was then aliquoted and frozen at -20°C for 2 months or until use, whichever occurred first. All dopamine preparations and treatments were performed in the dark using 10^-6^M dopamine unless otherwise stated, with dopamine treatment occurring immediately after thawing the dopamine. This concentration of dopamine was chosen to model the amount of dopamine to which macrophages in the CNS could be exposed to during stimulant use (Matt and Gaskill 2019). Although dopamine has been shown to oxidize and form cytotoxic reactive oxygen species in some *in vitro* conditions (Berman and Hastings 1999, Meiser, Weindl et al. 2013), previous data shows it is not associated with IL-1β production (Nolan, Reeb et al. 2020) and in our current studies we show that 10^-6^M dopamine is not cytotoxic to macrophages in our culture system (**Supplementary Figure 1**).

### 2.2 Isolation of primary peripheral blood mononuclear cells, monocytes, and generation of macrophages from human donors

Human peripheral blood mononuclear cells (PBMC) were separated from blood obtained from de-identified healthy donors from multiple sites including the New York Blood Center (Long Island City, NY, USA), University of Pennsylvania Human Immunology Core (Philadelphia, PA, USA), BioIVT (Westbury, NY, USA), and the Comprehensive NeuroHIV Center (CNHC) and Clinical and Translational Research Support Core (CNHC/CTRSC) Cohort at Drexel University College of Medicine (Philadelphia, PA, USA)) by Ficoll-Paque (GE Healthcare, Piscataway, NJ, USA) gradient centrifugation. PBMC were isolated and 5 million were set aside for subsequent mRNA analysis. Another 10 million PBMC were used to obtain pure, unlabeled monocytes by negative selection using the human Pan Monocyte Isolation kit (Miltenyi). Monocytes were set aside for subsequent qPCR. The remaining PBMC were matured into monocyte-derived macrophages (hMDM) using adherence isolation. Cells were cultured for 6-7 days in macrophage media (RPMI-1640 with 10% FBS, 5% human AB serum, 10 mM HEPES, 1% P/S, and 10 ng/mL M-CSF). A subset of cells from **Figure 5** were cultured in DMEM with 10% FBS, 5% human AB serum, 10 mM HEPES, 1% P/S, and 10 ng/mL M-CSF, and there was no significant difference relative to DMEM cultured samples (data not shown).

Limited, de-identified demographic information (age, gender, ethnicity, blood type and cytomegalovirus (CMV) status) was obtained as we have previously described (Matt, Nickoloff-Bybel et al. 2021), although all data categories were not available for each donor. The entire data set of donors was used to determine the relative expression of dopamine receptors, but not all demographic information was disclosed for every donor so not every donor was able to be used for every correlation. Dopamine receptor expression from subsets of these donors have been previously published (Nolan, Muir et al. 2018, Nickoloff-Bybel, Mackie et al. 2019, Matt, Nickoloff-Bybel et al. 2021), but this study combines donors from previous studies as well as 46 new donors across the current experiments. Use of the entire data set enables more robust examination of the correlations between IL-1β production and dopamine receptor expression due to the variability inherent in primary human macrophages.

### 2.3 Culture of C06 and C20 microglial cells

The C06 and C20 human microglial cell lines were a generous gift from the Karn laboratory (Case Western Reserve University) and details pertaining to the generation of these cell lines has been previously described (Garcia-Mesa, Jay et al. 2017). Briefly, human microglia were obtained from ScienCell Research Laboratories, Carlsbad, CA (Cat# HM1900) and then immortalized using simian virus 40 large T antigen and hTERT (Garcia-Mesa, Jay et al. 2017). These cells were maintained in 150-cm^2^ tissue culture flasks (Falcon) in DMEM supplemented with 5% FBS, 10 mM HEPES, 1% P/S, and 1% sodium pyruvate at 37°C in a humidified incubator under 5% CO_2_.

### 2.4 Differentiation and culture of human, iPSC-derived microglia

The inducible pluripotent stem cell-derived microglia (iMicroglia) were generated from common myeloid progenitors obtained from the Human Pluripotent Stem Cell Core at the Children’s Hospital of Philadelphia (CHOP) as we have done previously (Matt, Nickoloff-Bybel et al. 2021). This process uses an 11-day differentiation protocol that produces ramified cells that are susceptible to HIV infection and express the microglial markers CX_3_CR_1_, IBA1, TMEM119, and P2RY12, with very similar gene expression to human microglia (Ryan, Gonzalez et al. 2020). The cells used in this study were derived from the WT6 iPSC cell line. These cells were differentiated and maintained in RPMI-1640 supplemented with 1% FBS, 0.1% P/S, and IL-34 (100 ng/mL), M-CSF (25 ng/mL), and TGF-β1 (50 ng/mL) at 37°C in a humidified incubator under 5% CO_2_. Cytokines were added fresh with each media change. iMicroglia were cultured in glass bottomed petri dishes (Fluorodish, World Precision Instruments) or Cellbind plates (6 or 96 well, Fisher Scientific) depending on the experiment.

### 2.5 Lentiviral production and generation of C20.TetR.DRD2 microglial cell lines

The TetR and DRD2 lentivirus was made via calcium phosphate transduction of LentiX cells. TetR and helper plasmids and cell lines were a generous gift from Dr. Edward Hartsough (Drexel). Briefly, 3mg/mL of the PLP1, PLP2, PLPVSVG and TR (for TetR lines) or DRD2 (for DRD2 lines) plasmids were mixed with Ca^2+^. The solution was then added dropwise to 2X HBS and the resulting mix was added to the plate of LentiX cells at 70 – 80% confluency. After 4 days, supernatant was collected and incubated with 4X PEG-It solution for 24 hours. The supernatant-PEG-It mix was then spun down and the resulting viral pellet was resuspended in 150mL PBS and stored at -80°C until further use.

Wild-type C20 cells were transduced with 20mL concentrated TetR lentivirus for 48 hours. Following transduction, the cells were re-plated in 10cm dishes and treated with 8mg/mL blasticidin for 4 days. The concentration was then dropped to 5mg/mL blasticidin and media was refreshed every 3 days for approximately 3 weeks, until distinct colonies had formed. Individual colonies were picked and plated in individual wells of a 12-well plate and grown in C20 media without selection antibiotics until confluent. The cells were then re-plated in a 75cm^2^ flask and expanded. Successful transduction of 6 distinct colonies was confirmed using Western blotting. The clone with the highest expression, labeled C20.TetR #2, was used moving forward. The C20.TetR #2 cells were plated and transduced with DRD2 lentivirus at 20mL for 72 hours, moved to 10cm dishes and then selected for using in selection media (C20 media plus 300mg/mL Zeocin) for approximately 4 weeks as described above. Colonies were then picked and expanded until there were enough cells to be frozen down and stored for further experiments.

### 2.5 Viral stocks

Viral stocks of HIV_ADA_ were generated by infecting CEM-SS cells with HIV_ADA_, a blood derived, R5-tropic strain of HIV (Gendelman, Orenstein et al. 1988), as we have done previously (Gaskill, Calderon et al. 2009, Matt, Nickoloff-Bybel et al. 2021). Briefly, CEM-SS cells cultured in T175 flasks were infected with a single tube of HIV_ADA_ stock, then cultured until syncytia started to form, changing media every 3 – 4 days. For the stocks used in this study, syncytia began to form at 17 days post infection, so cell-free supernatants were collected daily at 18 to 41 days post-infection. Each stock was centrifuged at 1200 x g to pellet cells and debris, then moved to a fresh conical vial, aliquoted in 1 mL aliquots and stored at -80°C for use as viral stocks. Stock concentration was determined by quantifying the amount of p24•Gag per mL, using an HIV p24 (high sensitivity) AlphaLISA Detection kit (Perkin-Elmer, Waltham, MA).

### 2.6 Cell viability assays

Primary hMDM were plated at 15,000 cells/well in a 96-well black-walled plate (NUNC, ThermoFisher Scientific) and treated with a range of concentrations of dopamine (1×10^-10^M to 5×10^-4^M) in duplicate wells for 24 hours. After treatment, supernatants were collected, and cells were stained with LIVE/DEAD Cell Imaging Kit (ThermoFisher Scientific cat # R37601) according to manufacturer’s instructions. Cells were then imaged using the Cellomics CellInsight CX7 HCS Platform at 20x magnification and 20 fields per well. After images were acquired on the CX7, quantitative analysis was performed, to identify live and dead macrophages using HCS Studio software, using the Thermo Scientific Cellomics Colocalization Bioapplication, and specific parameters are included in **Supplementary Table 1**. These images had donor-specific manual alterations in Image Settings to correct any background. As an example for one representative donor, the live (green) stain was gated by excluding border objects, thresholding (fixed) at 500, segmentation (intensity) at 500, object area 854.84-7746.16 to eliminate small non-cellular debris, object variant intensity 7.52 – 65535 (**Supplementary Table 1**). The dead (red) stain was gated by excluding border objects, thresholding (fixed) at 2000, segmentation (intensity) 28, and object area 265.28 to 6653.4 to exclude small debris. Cell level data was taken at Object Count Ch2 and used to calculate total number of cells, live cells, dead cells and %viability. To examine the initial health of the hMDM cultures pre-experiment, a time 0 was added to a subset of donors.

To provide a confirmatory evaluation of cell viability, collected supernatants were quantified for LDH release, using the CyQUANT^tm^ LDH Cytotoxicity Assay (ThermoFisher Scientific # C20301). Supernatants from time 0 were used as a spontaneous LDH release control to quantify change in LDH over 24 hours in vehicle and treated wells. Cellular lysates using M-PER (ThermoFisher Scientific # PI78501) were used to identify the maximum LDH release possible. 50 μL of each condition were loaded in triplicate wells on 96-well plates and incubated with 50 μL of reaction mixture (LDH substrate mix, and assay buffer) for 30 minutes at 25°C in the dark. Stop solution (50mL) was added to each well and then absorbance values at 490nm and 680nm were taken on a spectrophotometer. Viability was calculated by first calculating the % cytotoxicity using the equation 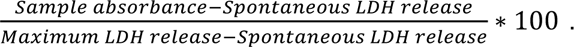 This percentage was then subtracted from 100% to obtain the %viability.

### 2.7 Quantitative RT-PCR

Total RNA was extracted from cells using Trizol, the RNeasy Mini Plus^™^ kit (Qiagen), or the *Quick*-DNA/RNA Microprep Plus Kit (Zymo), and RNA quantity and purity were determined by a NanoDropOne spectrophotometer (Nanodrop Technologies). RNA (1 μg) was used to synthesize cDNA using the high-capacity reverse transcriptase cDNA synthesis kit (Abcam). All dopamine receptor subtypes (DRD1-DRD5), IL-1β, NLRP3, NLRP1, NLRC5, NLRC4, AIM2, IL-18, IL-1rn, PYCARD, CASP1, and the housekeeping gene 18s were amplified from cDNA by quantitative PCR (qPCR) on a QuantStudio 7 using gene-specific primers.

### 2.8 RNA-seq library preparation

RNA-seq library preparation was performed as described elsewhere and as follows (Link, Duttke et al. 2018). RNA isolation from C20 cells was carried out as described above and total RNA was enriched in poly-A tailed RNA transcripts by double incubation with Oligo d(T) Magnetic Beads (NEB, S1419S) and fragmented for 9 minutes at 94°C in 2X Superscript III first-strand buffer containing 10 mM DTT (Invitrogen, #P2325). The 10 μL of fragmented RNA was added to 0.5 μL of Random primers (Invitrogen, #48190011), 0.5 μL of Oligo d(T) primer (Invitrogen, #18418020), 0.5 μL of SUPERase inhibitor (Ambion, #AM2696), 1 μL of 10 mM dNTPs and incubated at 50°C for 3 minutes. Then, 5.8 μL of water, 1 μL of 10 mM DTT, 0.1 μL of 2 μg/μL Actinomycin D (Sigma, #A1410), 0.2 μL of 1% Tween-20 (Sigma) and 0.5 μL of SuperScript III (Invitrogen, #ThermoFisher 18080044) was added to the mix. Reverse-transcription (RT) reaction was performed at 25°C for 10 minutes followed by 50°C for 50 minutes. RT product was purified with RNAClean XP (Beckman Coulter, #A63987) and eluted in 10 μL in 0.01% Tween 20. The RNA-cDNA complex was then added to 1.5 μL of 10X Blue Buffer (Enzymatics, #B0110-L), 1.1 μL of dUTP mix (10 mM dATP, dCTP, dGTP and 20 mM dUTP), 0.2 μL of 5 μ/mL RNAseH (Enzymatics, #Y9220L), 1 μL of 10 U/μl DNA polymerase I (Enzymatics, #P7050L), 0.15 μL of 1% Tween 20 and 1.05 μL of nuclease free water; and incubated at 16°C for 2.5 hours or overnight. The resulting dsDNA product was purified using 28 μL of Sera-Mag SpeedBeads Magnetic Carboxylate (ThermoFisher, #65152105050250) diluted in 20% PEG8000, 2.5 M NaCl to a final 13% PEG concentration, washed twice with 80% EtOH, air dry and eluted in 40 μL of 0.05% Tween 20. The purified 40 μL of dsDNA was end-repaired by blunting followed by A-tailing and adapter ligation as described elsewhere (Heinz, Benner et al. 2010) using BIOO Barcodes (BIOO Scientific, #514104), 0.33 μL 1% Tween 20 and 0.5 μL T4 DNA ligase HC (Enzymatics, #L6030-HC-L). Libraries were amplified by PCR for 11–15 cycles using Solexa IGA and Solexa IGB primers (AATGATACGGCGACCACCGA and CAAGCAGAAGACGGCATACGA, respectively), and size selected for fragments (200-500 bp) by gel extraction (10% TBE gels, Life Technologies EC62752BOX). Eluted libraries were quantified using a Qubit dsDNA HS Assay Kit and sequenced on a NovaSeq 6000 (Illumina, San Diego, California).

### 2.9 Analysis of RNA-Seq

To generate the read count matrix, we used nf-core RNAseq pipeline version 3.10.1 with the FASTQ sequencing files as input (Di Tommaso, Chatzou et al. 2017, Harshil Patel 2023). The reads are filtered and mapped to a customized reference genome which combines hg38 and NC_001802.1 using STAR aligner (Dobin, Davis et al. 2013). The resulting count matrix is used as the input for Partek® Flow® software for differential expression analysis. We used DESeq2 to conduct the differential expression analysis, and the results are filtered with FDR < 0.05 (Love, Huber et al. 2014). For functional analysis, we used IPA on the filtered gene lists (QIAGEN Inc., https://digitalinsights.qiagen.com/IPA).

### 2.10 IL-1β production and caspase-1 secretion assays

Primary hMDM were cultured at 9.5 x 10^4^ cells per well in 48-well plates (BD Falcon), C06/C20 microglia were cultured at 1 x 10^6^ cells per well in 6-well plates (BD Falcon), and iMicroglia were cultured at 5 x 10^4^ cells per well in 96-well plates (CellBind). Macrophages (hMDM) were treated for 24 hours (Nolan, Muir et al. 2018) and microglia (C06/C20 and iMicroglia) were treated for 4 hours with 10^−6^M dopamine. Treatment with LPS (10 ng/mL) was used as a positive control. At 30 minutes (hMDM) or 1 hour (C06/C20 and iMicroglia) prior to supernatant collection, cells were stimulated with ATP (2.5 mM) to induce inflammasome activation, and lysates were collected for the analysis of IL-1β production while supernatants were collected for the analysis of active caspase-1. Lysates were analyzed for IL-1β using AlphaLISAs performed according to the manufacturer’s protocol (PerkinElmer). The limit of detection for the IL-1β AlphaLISAs was 0.6 pg/mL. Cells that did not respond to LPS with at least a 1.5-fold increase in IL-1β were excluded from the analysis. The caspase-1 concentration in supernatants was determined via Quantikine® ELISA performed according to the manufacturer’s protocol, and the limit of detection for this assay was 0.68 pg/mL (R&D systems, Minneapolis, MN, USA).

### 2.11 Immunofluorescence

For immunofluorescent staining of microglial markers, C20 cells were cultured in 96-well glass-like polymer plates (P96-1.5P, Cellvis) at a density of 5,000 cells per well. Cells reached optimal confluency after 4 days, at which point cells were fixed with 4% PFA at room temperature for 10 minutes, permeabilized, and incubated with blocking buffer (1% BSA, 0.1% Tween 20, and 22.52 mg/mL glycine in 1X PBS) at room temperature for 30 minutes. After blocking, cells were then incubated overnight at 4°C with a P2RY12 primary antibody (1:100, 702516, Thermo Fisher) or TMEM119 primary antibody (1:50, PA5119902, Thermo Fisher) diluted in blocking buffer. Following the primary incubation, the C20 cells were incubated with an Alexa Fluor 488 secondary antibody (A-11001, Fisher Scientific) diluted in blocking buffer for 1 hour at room temperature. All cells were stained with DAPI (0.2 µg/mL, D1306, Thermo Fisher) for 10 minutes and preserved in 1X PBS. High-content imaging of stained cells was performed using the Cellomics CX7 platform, capturing 20 fields per well at 20x magnification.

To prepare iMicroglia for immunofluorescent staining, cells were cultured in glass-bottomed petri dishes (Fluodish, World Precision Instruments) coated with poly-D-lysine (0.1 mg/mL, P6407, Sigma-Aldrich) at a density of 300,000 cells per dish. The cells were either mock-infected or infected with HIV, in the presence or absence of 10^-6^M dopamine, for 7 days. After the treatment, the cells were fixed with 4% PFA at room temperature for 10 minutes, permeabilized, and incubated with blocking buffer (1% BSA, 0.1% Tween 20, and 22.52 mg/mL glycine in 1X PBS) at room temperature for 30 minutes. Subsequently, the cells were incubated overnight at 4°C with an anti-HIV-1 p24 monoclonal (AG3.0) primary antibody (4 µl/mL) (NIH HIV Reagent Program catalog# 4121) or P2RY12 primary antibody (1:100, 702516, Thermo Fisher) or TMEM119 primary antibody (1:50, PA5119902, Thermo Fisher) or Iba1 (10ug/mL, PA5121836, Thermo Fisher) prepared in blocking buffer. After the primary incubation, iMicroglia were incubated with an Alexa Fluor 488 secondary antibody (A-11001, Fisher Scientific) prepared in blocking buffer for 1 hour at room temperature. Afterwards, all cells were stained with CellMask Deep Red (250ng/mL, catalog # C10046 Thermo Fisher) for 10 minutes and cover slipped with SlowFade™ Glass Soft-set Antifade Mountant, with DAPI (catalog # S36920 Thermo Fisher). iMicroglia images were acquired at 60X using an Olympus FV3000 confocal laser scanning microscope.

### 2.12 Western blotting

For Western blot analysis, hMDM were cultured at 9.5 x 10^5^ cells per well in 6-well plates (Thermo Fisher). The cells were mock-infected or infected with HIV for 7 days, and after 7 days of infection, treated with vehicle (H_2_O) or 10^-6^M dopamine for 3 hours. Cells were lysed using lysis buffer from the Quick-DNA/RNA Microprep Plus Kit (D7005, Zymo Research), and flow-through protein was separated according to the manufacturer’s instructions. Cold acetone was added to the flow-through protein in a 4:1 ratio and the mixture was incubated on ice for 30 minutes. Then samples were centrifuged at 13,000 RPM for 10 minutes at 4°C. Following aspiration of the supernatant, samples were washed in 400 μl of molecular-grade 100% EtOH. The samples were again centrifuged at 13,000 RPM for 1 minute at 4°C, supernatants were removed and then protein pellets were dissolved in mammalian protein extraction reagent (M-PER, Thermo Fisher Scientific, Waltham, MA), containing 1% Halt Protease and Phosphatase Inhibitor cocktail and 1% EDTA (Thermo Fisher Scientific, Waltham, MA). Lysates were sonicated with a Q125 sonicator (Qsonica, Newtown, CT) at 25% power for 5 seconds and stored at 4°C for 1 – 7 days. Subsequently, protein concentrations were quantified using a Bicinchoninic acid assay (BCA) with the Pierce BCA Protein Assay Kit (Thermo Fisher Scientific). The lysates were diluted to a concentration of 1.5 μg/μL and stored at -80°C until analyzed by Western blot.

On the day of the Western blot, samples were thawed at RT, and mixed with 4X protein loading buffer (Licor Biosciences, Lincoln, NE) containing 10% 2-Mercaptoethanol (BME), then boiled at 95°C for 10 minutes. Protein lysates were then separated by gel electrophoresis on Bolt Bis-Tris Plus gradient 4-12% precast gels in MOPS/SDS running buffer in a Mini gel tank (Life Technologies, Carlsbad CA). Separation was performed for 60 minutes at 150V, and then proteins were transferred to an Immobilon PVDF membrane (EMD Millipore, Temecula, CA) at 25V for 90 minutes. To generate an internal loading control, membranes were imaged after treatment with Revert Total Protein Stain (LI-COR Biosciences, Lincoln, NE) according to the manufacturer’s instructions. Revert stain was then removed and then membranes were blocked in 5% BSA for 2 hours at RT and then incubated overnight at 4°C in either anti-NLRP3 antibody (15101S, 1:1000 in 5% BSA, Cell Signaling), anti-NLRC4 antibody (12421, 1:1000 in 5% BSA, Cell Signaling) or anti-AIM2 antibody (MA5-38442, 1:1000 in 5% BSA, Thermo Fisher Scientific). Following primary antibody incubation overnight, blots were washed in TBS with 0.1% Tween, stained with anti-rabbit IgG HRP linked antibody (CST 7074, 1:3000 in 5% milk) for NLRP3 and NLRC4 or anti-mouse IgG HRP linked antibody (CST 7076, 1:3000 in 5% milk) for AIM2, and incubated at RT for 1 hour. After the secondary incubation, blots were washed and incubated in Supersignal West Pico PLUS plus Chemiluminescent Substrate (2 mL, 30 sec, ThermoFisher, 34580). Blots were imaged using an Odyssey Fc Imaging System and analyzed using Image Studio Lite (Licor Biosciences, Lincoln, NE). Target bands were normalized to the total protein stain, and then each condition was compared to the mean expression of the vehicle controls to determine the fold change in expression.

### 2.13 Statistics

Prior to analysis, all data were normalized to the mean of the vehicle treated condition. To determine the appropriate statistical tests, data sets were evaluated by analysis of skewness and evaluation of normality and lognormality to determine the distribution of the data. Extreme data points presumed to be technical outliers were identified via ROUT test (Q = 0.1%) and removed from analysis. Statistical analysis of gene expression data was performed on data normalized to 2^-ΔC^_T_. *Post-hoc* analyses were performed when appropriate. While positive controls are shown on the same graph as dopamine-mediated changes, they were not included in ANOVA analyses analyzing the impact of dopamine in most figures. The separate tests performed on the positive controls are denoted using the @ sign, rather than the * used to show significance in the analyses of dopamine-mediated changes. All data analysis was performed using GraphPad Prism 10.0 (Graphpad, La Jolla, CA). p < 0.05 was considered significant.

### 2.14 Data Availability

The RNA-seq datasets in the current study will be deposited in GEO.

## 3. Results

### 3.1 Dopamine-mediated increase in IL-1β production in primary macrophages varies with expression of dopamine receptor subtypes

Our prior work on dopamine-mediated regulation of cytokine production in primary human monocyte-derived macrophages (hMDM) (Nolan, Muir et al. 2018, Nolan, Reeb et al. 2020) suggests a pivotal role of dopamine receptor subtype expression on cytokine production. Across hMDM examined from 42 healthy donors, we observed a significant increase in IL-1β protein production in response to 10^-6^M dopamine (**Figure 1A**, Wilcoxon test, n = 42, * p = 0.0276, sum of (+,-) ranks 627, -276). Next, we determined the expression of dopamine receptors in hMDM via qPCR. From these 42 donors, most hMDM express DRD1 (71.43%) and DRD2 (64.29%), whereas DRD3 (21.43%) and DRD4 (30.95%) are expressed on a much smaller proportion of hMDM (**Figure 1B**). DRD5 mRNA was detected in all samples.

**Figure 1.**
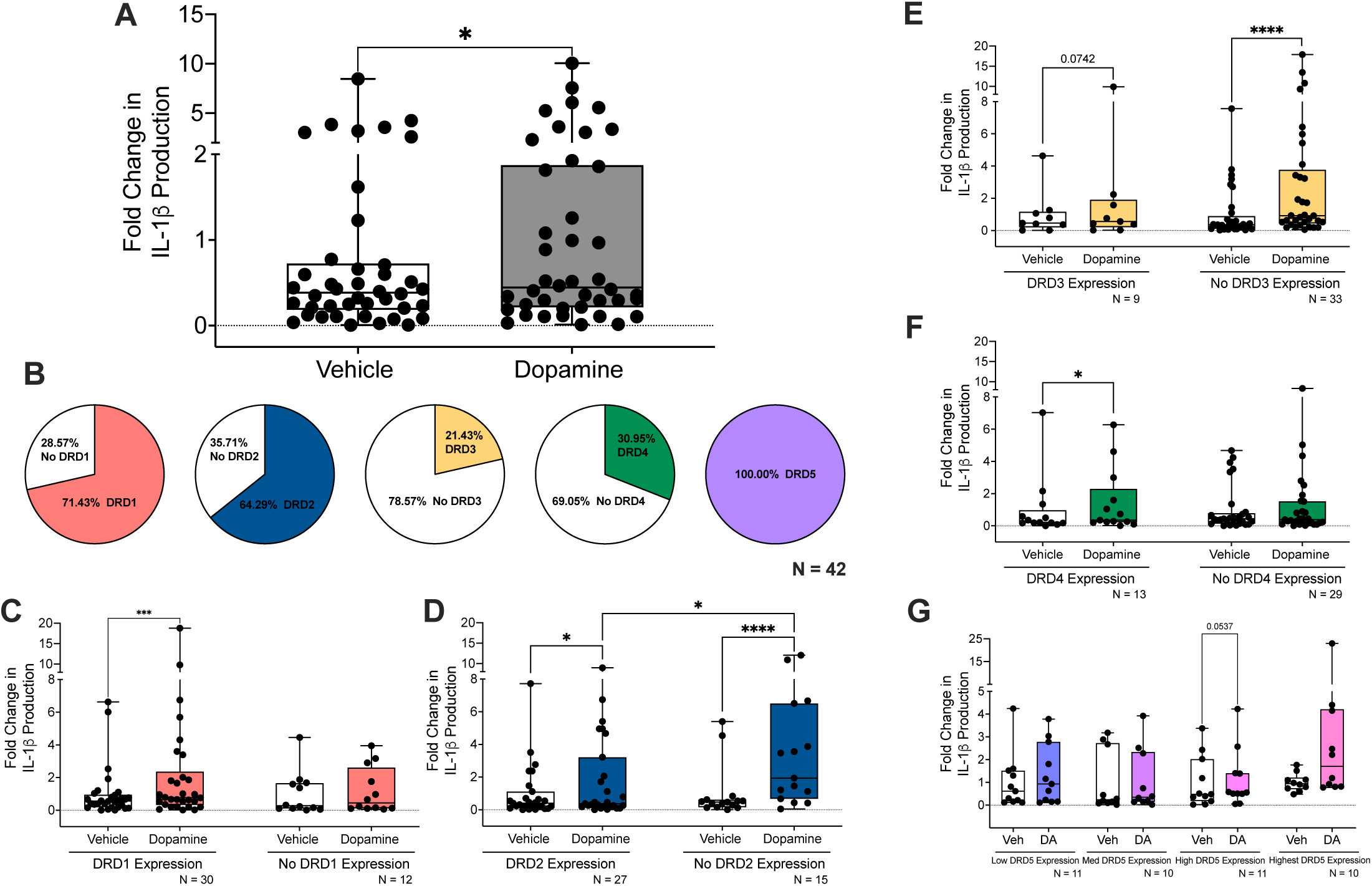
Dopamine-mediated increase in IL-1β production in primary macrophages varies with expression of dopamine receptor subtypes. Primary human monocyte-derived macrophages (hMDM) were treated for 24 h with 10^-6^ M dopamine. Lysates were collected and analyzed for IL-1β using AlphaLISAs. Results were normalized to protein concentration. qPCR detected mRNA for all subtypes of dopamine receptors (DRD1, DRD2, DRD3, DRD4 and DRD5). In our 42 donors, which together significantly respond to dopamine and increase IL-1β production **(A)**, most express DRD1 and DRD2, whereas a much smaller proportion express DRD3 and DRD4 **(B)**. The connection between dopamine receptors and IL-1β was determined by analysis of IL-1β levels in hMDM that did or did not express **(C)** DRD1, **(D)** DRD2, **(E)** DRD3, and **(F)** DRD4. Although all express DRD5, we examined DRD5’s impact on dopamine-induced IL-1β production by splitting expression into quartiles: low, medium, high, and highest expression **(G)**. Significance was determined using Wilcoxon and Mann-Whitney tests, *p < 0.05, ***p < 0.001, and ****p < 0.0001.

Next, we wondered if differential dopamine receptor expression impacts IL-1β production. We detected a significant dopamine-mediated increase in IL-1β production in hMDM with detectable DRD1 expression compared to samples with non-detectable DRD1 mRNA (**Figure 1C**, Wilcoxon tests, with DRD1, n = 30, *** p = 0.0009, sum of (+,-) ranks 388, -77; without DRD1, n = 12, p = 0.4697, sum of (+,-) ranks 49, -29). In contrast, dopamine significantly increased IL-1β production in hMDM both with and without DRD2 detectable expression. However, in hMDM without DRD2, the magnitude of the dopamine-mediated increase was significantly greater than in hMDM with DRD2 (**Figure 1D**, Wilcoxon tests, with DRD2, n = 27, * p = 0.0260, sum of (+,-) ranks 281, -97; without DRD2, n = 15, **** p < 0.0001, sum of (+,-) ranks 120, 0; Mann-Whitney test, n = 15 - 27, *p = 0.0277, sum of (Dopamine-induced IL-1β D1 vs No D1) ranks 497, 406, U=119). The effects of DRD3 expression were similar to those found with DRD2, with dopamine-mediated increases in IL-1β in hMDM both with and without detectable DRD3, although this was only significant in hMDM without DRD3 (**Figure 1E**, Wilcoxon tests, with DRD3, n = 9, p = 0.0742, sum of (+,-) ranks 38, -7; without DRD3, n = 33, ****p < 0.0001, sum of (+,-) ranks 537, -24). Dopamine-mediated IL-1β production was dependent on detectable DRD4 expression, like DRD1 (**Figure 1F**, Wilcoxon tests, with DRD4, n = 13, *p = 0.0398, sum of (+,-) ranks 75, -16; without DRD4, n = 29, p = 0.8148, sum of (+,-) ranks 229, -206). Finally, while all hMDM we examined express DRD5, we wanted to determine if different levels of DRD5 expression might affect dopamine-induced IL-1β production. To do this, we split donor DRD5 expression into quartiles and examined the effect of dopamine in each group. While there was a trend toward a significantly increased impact of dopamine on IL-1β production in the 3^rd^ quartile, no other effect of differing DRD5 expression was discerned (**Figure 1G**, Wilcoxon tests, low DRD5, n = 11, p = 0.1748, sum of (+,-) ranks 49, -17; medium DRD5, n = 10, p = 0.6953, sum of (+,-) ranks 32, -23, high DRD5, n = 11, p = 0.0537, sum of (+,-) ranks 55, -1, highest DRD5, n = 10, p = 0.1309, sum of (+,-) ranks 43, -12). Taken together, these data indicate that expression of both D1-like and D2-like receptors can influence IL-1β production in human macrophages.

Recent findings indicate that the relative expression of D1-like vs. D2-like dopamine receptors (i.e. receptor ratio) shapes excitatory versus inhibitory signaling, potentially playing a role in subsequent neuropathologies and cognitive dysfunction (Mandeville, Sander et al. 2013, Manza, Shokri-Kojori et al. 2022). Indeed, dopamine receptor ratios are diverse across brain regions, change with substances of misuse, and can influence drug seeking behavior (Thompson, Martini et al. 2010, Sillivan and Konradi 2011). It has also been suggested that dopamine receptor ratios could be equally important in immune cells and the regulation of inflammatory signaling (Pacheco, Contreras et al. 2014). Therefore, we further stratified our data to examine how the combination of expression or absence of two dopamine receptors might influence dopamine-mediated increases in IL-1β (**Table 1**). The only receptor combination where we detected significant dopamine-mediated increases in IL-1β production was in samples with detectable DRD3 and DRD4 expression (Wilcoxon tests, DRD3+ DRD4+, n = 7, *p = 0.0156, sum of (+,-) ranks 28, 0; DRD3-DRD4+, n = 6, p = 0.5625, sum of (+,-) ranks 14, -7, DRD3+ DRD4-, n = 2, N/A, DRD3-DRD4-, n = 27, p = 0.1482, sum of (+,-) ranks 250, -128). These data suggest that the specific combination of dopamine receptors expressed may be just as important as expression of a single receptor when mediating dopamine’s effects on IL-1β.

**Table 1:** Different combinations of dopamine receptors expressed on primary macrophages changes dopamine’s effects on IL-1b production.

| Stratification | N | Dopamine-mediated Fold Change | p value | SD |
| --- | --- | --- | --- | --- |
| DRD1+ DRD2+ | 19 | 2.36 | 0.087 | 4.09 |
| DRD1- DRD2+ | 8 | 1.48 | 0.188 | 1.39 |
| DRD1+ DRD2- | 11 | 1.26 | 0.278 | 1.30 |
| DRD1- DRD2- | 4 | 1.34 | 0.625 | 1.43 |
| DRD1+ DRD3+ | 4 | 1.37 | 0.125 | 1.70 |
| DRD1- DRD3+ | 5 | 1.86 | 0.313 | 2.82 |
| DRD1+ DRD3- | 26 | 1.95 | 0.075 | 3.11 |
| DRD1- DRD3- | 7 | 1.05 | 0.938 | 1.12 |
| DRD1+ DRD4+ | 8 | 2.14 | 0.055 | 2.78 |
| DRD1- DRD4+ | 5 | 1.23 | 0.438 | 1.39 |
| DRD1+ DRD4- | 22 | 1.87 | 0.166 | 3.24 |
| DRD1- DRD4- | 7 | 1.15 | 0.938 | 1.47 |
| DRD2+ DRD3+ | 7 | 1.78 | 0.156 | 2.84 |
| DRD2- DRD3+ | 2 | 1.39 | N/A | 1.79 |
| DRD2+ DRD3- | 20 | 1.59 | 0.294 | 2.31 |
| DRD2- DRD3- | 13 | 1.28 | 0.244 | 1.22 |
| DRD2+ DRD4+ | 9 | 1.46 | 0.055 | 1.58 |
| DRD2- DRD4+ | 4 | 1.02 | 0.875 | 0.82 |
| DRD2+ DRD4- | 18 | 1.78 | 0.640 | 3.26 |
| DRD2- DRD4- | 11 | 1.39 | 0.169 | 1.05 |
| DRD3+ DRD4+ | 7 | 1.89 | 0.016 | 2.96 |
| DRD3- DRD4+ | 6 | 1.25 | 0.563 | 1.58 |
| DRD3+ DRD4- | 2 | 0.51 | N/A | 0.65 |
| DRD3- DRD4- | 27 | 1.60 | 0.148 | 2.41 |

We also examined whether demographic details including sex, CMV status, and age influenced the observed dopamine-mediated increase in IL-1β production. Understanding these relationships is important as regulation of sex hormones can impact dopamine synthesis, release, turnover, and degradation (Becker 1990, Watson, Alyea et al. 2006, Sinclair, Purves-Tyson et al. 2014), CMV virus can alter catecholamine metabolism (O’Kusky, Boyes et al. 1991) which could contribute to risk for psychiatric disorders (Burgdorf, Trabjerg et al. 2019), and dopaminergic dysfunction plays a role in modulating oxidative stress and inflammation in the elderly (Villar-Cheda, Dominguez-Meijide et al. 2014, Garrido-Gil, Dominguez-Meijide et al. 2018). Interestingly, while hMDM from males showed a significant dopamine-mediated increase in IL-1β production, the magnitude of the dopamine-induced fold change in IL-1β was greatest in hMDM from females (**Supplementary Figure 2A**, Wilcoxon tests, Male, n = 23, *p = 0.0354, sum of (+,-) ranks 207, -69; Female, n = 17, p = 0.7119, sum of (+,-) ranks 85, -68). As expression of different dopamine receptor subtypes might influence the effect of dopamine on IL-1β, we then examined the distribution of dopamine receptor subtypes on hMDM from male and female donors. While there were small differences in the expression of the receptors in hMDM from males and females, there was no statistical difference in expression of any of the dopamine receptor subtypes (**Supplementary Figure 2B**). We also found that infection with CMV, which is common in the adult population (Griffiths and Reeves 2021, Fowler, Mucha et al. 2022), does not significantly impact dopamine-mediated changes in IL-1β production (**Supplementary Figure 2C**, Wilcoxon tests, CMV-, n = 12, p = 0.3013, sum of (+,-) ranks 53, -25; CMV+, n = 15, p = 0.3591, sum of (+,-) ranks 77, -43). This was somewhat surprising, as our prior publication (Nolan, Reeb et al. 2020) showed that in CMV-infected donors, dopamine increased NF-κB activity, a precursor to IL-1β production. However, the dopamine-induced fold change in IL-1β was greatest in donors who were CMV+ (fold change of 2.186 +/- 0.8615 SEM (CMV+) vs. fold change of 1.388 +/- 0.5214 SEM (CMV-)). Finally, age-related analyses showed a trend in that donors under 50 had a dopamine-mediated increase in IL-1β production, although the dopamine-induced fold change in IL-1β was greatest in donors over 50 (**Supplementary Figure 2D**, Wilcoxon tests, Under 50, n = 29, p = 0.0724, sum of (+,-) ranks 301, -134; Over 50, n = 11, p = 0.5195, sum of (+,-) ranks 41, -25). These data suggest that a proportion of the heterogeneity we see with dopamine-mediated changes in IL-1β may be attributable to demographic factors.

As IL-1β production can be influenced by ATP and other damage associated molecular patterns associated with cell death, we evaluated the cytotoxicity of the dopamine concentrations used in these studies. A subset of donors was treated with a range of dopamine concentrations for 24 hours and assessed using a LIVE/DEAD cell imaging assay (n = 5) or quantification of LDH release (n = 8). In both assays, 24 hours of vehicle treatment showed a 2 – 3% decrease in viability. Compared with vehicle treated cells, neither assay showed a significant decrease in viability in response to 10^-6^M dopamine, as used in our study. However, the two highest concentrations of dopamine used (10^-4^M and 5×10^-4^M) did show a slight but significant decrease in cell viability, from 97.927% in vehicle to 92.549% at 10^-4^M and 86.028% at 5×10^-4^M. Representative images of the LIVE/DEAD assay are shown (**Supplementary Figure 1A**) along with the pooled data (**Supplementary Figure 1B**, Friedman test, n = 5, Friedman statistic 24.37, **p = 0.020; *Post-hoc* with Dunn’s multiple comparisons, vehicle vs. 1×10^-4^M: *p = 0.0214, vehicle vs. 5×10^-4^M **p = 0.0065). Analysis of cell viability by LDH assay showed similar results, with the highest three concentrations of dopamine reducing viability from 96.15% in vehicle to 89.4% at 10^-5^M, 86% at 10^-4^M and 80.5% at 5×10^-4^M (**Supplementary Figure 1C**, Friedman test, n = 8, Friedman statistic 54.33, **** p <0.0001; P*ost-hoc* with Dunn’s multiple comparisons, vehicle vs. 1×10^-5^M, ***p = 0.001, vehicle vs. 1×10^-4^M ****p <0.0001, vehicle vs. 5×10^-4^M ****p <0.0001). These data show that while higher concentrations do produce some cytotoxicity, the concentration of dopamine used for our analyses (10^-6^M) produced no significant changes in hMDM viability.

### 3.2 IL-1β and dopamine receptor expression levels in primary human myeloid cells change with maturation

A number of studies have examined dopamine receptor gene and protein expression in myeloid cells (Mastroeni, Grover et al. 2009, Gaskill, Carvallo et al. 2012, Coley, Calderon et al. 2015, Huck, Freyer et al. 2015, Nolan, Muir et al. 2018, Nickoloff-Bybel, Mackie et al. 2019, Matt, Nickoloff-Bybel et al. 2021). However, it is not clear if the expression is a characteristic of cell type and/or if maturation from monocytes to macrophages could affect receptor levels. To examine this, we evaluated how dopamine receptor expression changed in PBMC, monocytes and hMDM, many of which were derived from the same donor. There was significantly higher D1-like receptor expression in mature hMDM than in PBMC and monocytes (**Figure 2A**, Kruskal-Wallis test, n = 33 – 42; **DRD1** Kruskal-Wallis statistic 16.48, ***p = 0.0003, *Post-hoc* with Dunn’s multiple comparisons, Monocytes vs. hMDM, ***p = 0.0008, hMDM vs PBMC, **p = 0.0021, Monocytes vs PBMC, p > 0.999; **DRD5**, Kruskal-Wallis statistic 19.41, ****p < 0.0001, *Post-hoc* with Dunn’s multiple comparisons, Monocytes vs. hMDM, **p = 0.0013, hMDM vs PBMC, ***p = 0.0002, Monocytes vs PBMC, p > 0.999). The D2-like receptors were more divergent, with only DRD2 showing significantly higher expression in hMDM compared to PBMC and monocytes (**Figure 2A**, Kruskal-Wallis test, n = 33 - 42, Kruskal-Wallis statistic 13.56, **p = 0.0011, *Post-hoc* with Dunn’s multiple comparisons, Monocytes vs. hMDM, **p = 0.0011, hMDM vs PBMC, **p = 0.0299, Monocytes vs PBMC, p = 0.7225). Expression of DRD3 was significantly higher expression in PBMC than in hMDM and monocytes (**Figure 2A**, Kruskal-Wallis test, n = 33 - 42, Kruskal-Wallis statistic 41.21, ****p < 0.0001, *Post-hoc* with Dunn’s multiple comparisons, Monocytes vs. hMDM, p = 0.8594, hMDM vs PBMC, ****p < 0.0001, Monocytes vs PBMC, ****p < 0.0001). Further, DRD4 expression was significantly lower in hMDM than in both PBMC and monocytes (**Figure 2A**, Kruskal-Wallis test, n = 33 - 42, Kruskal-Wallis statistic 77.21, ****p < 0.0001, *Post-hoc* with Dunn’s multiple comparisons, Monocytes vs. hMDM, ****p < 0.0001, hMDM vs PBMC, ****p < 0.0001, Monocytes vs PBMC, p = 0.0712).

**Figure 2.**
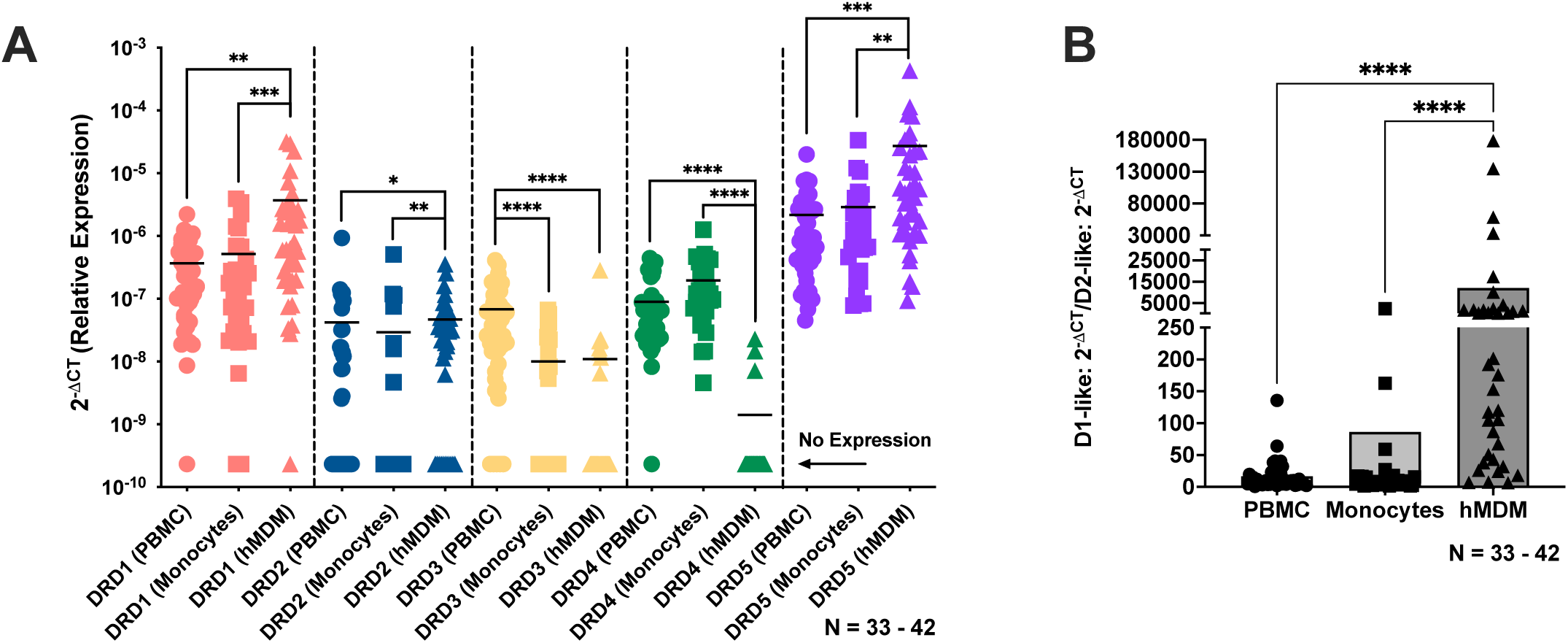
Dopamine receptor expression levels in primary human myeloid cells change with maturation. Primary human peripheral blood mononuclear cells (PBMCs), monocytes, and monocyte-derived macrophages (hMDM) were collected for subsequent mRNA analysis. 19 of these donors were paired across the three cell types. **(A)** qPCR detected mRNA for all subtypes of dopamine receptors (DRD1, DRD2, DRD3, DRD4 and DRD5) that were differentially expressed in these cell types (N = 33 - 42). **(B)** Using a D1-like to D2-like ratio (DRD1 + DRD5 / (DRD2 + DRD3 + DRD4), hMDM had a significantly higher ratio compared to monocytes or PBMCs. Significance was determined using Kruskal-Wallis tests and post-hocs with Dunn’s multiple comparisons,*p < 0.05, **p < 0.01, ***p < 0.001, and ****p < 0.0001.

Another way to examine changes in relative receptor expression is to determine the ratio of D1-like to D2-like receptors, by pooling the values for D1-like receptor expression and dividing them by the pooled values for D2-like expression for every donor, as we have done previously (Matt, Nickoloff-Bybel et al. 2021). Analysis of D1-like / D2-like ratios found that hMDM have a significantly higher and more variable D1-like to D2-like receptor ratio (12169 ± 37086) compared to monocytes (86.36 ± 411.9) or PBMCs (16.62 ± 23.5) (**Figure 2B**, Kruskal-Wallis test, n = 33 - 42, Kruskal-Wallis statistic 51.46, ****p < 0.0001, *Post-hoc* with Dunn’s multiple comparisons, Monocytes vs. hMDM, ****p < 0.0001, Monocytes vs PBMC, p > 0.999, hMDM vs PBMC, ****p < 0.0001). This shows that dopamine receptor expression levels in macrophages are distinct from those in monocytes and PBMCs. Using these data, we then correlated dopamine receptor mRNA expression with IL-1β mRNA expression in each cell population. Expression of DRD1, DRD2, DRD3, and DRD5 were positively correlated with IL-1β expression in hMDM (**Supplementary Figure 3A-D**, hMDM IL-1β vs DRD1, n = 28, Spearman r = 0.4921, **p = 0.0078, hMDM IL-1β vs DRD2, n = 28, Spearman r = 0.4910, *p = 0.0265, hMDM IL-1β vs DRD3, n = 28, Spearman r = 0.6134, ***p = 0.0005, hMDM IL-1β vs DRD5, n = 28, Spearman r = 0.4877, **p = 0.0085). In monocytes, the only significant correlation was a positive correlation between DRD2 and IL-1β (**Supplementary Figure 3E**, Monocytes IL-1β vs DRD2, n = 28, Spearman r = 0.4735, *p = 0.0109), and there were no positive correlations in PBMC.

The expression and release of IL-1β differs between myeloid cell types, including primary human monocytes and MDM (Netea, Nold-Petry et al. 2009, Hadadi, Zhang et al. 2016, Awad, Assrawi et al. 2017), as well as primary macaque monocytes, bone marrow-derived macrophages, and microglia (Burm, Zuiderwijk-Sick et al. 2015). To confirm that the observed correlations were not solely due to differences in baseline IL-1β, we examined levels of IL-1β transcript in untreated PBMC, monocytes and hMDM. Monocytes expressed significantly higher IL-1β mRNA relative to hMDM, but no other differences were observed between the three cell populations (**Supplementary Figure 3F**, Kruskal-Wallis test, n = 28 - 32, Kruskal-Wallis statistic 9.905, **p = 0.0071, *Post-hoc* with Dunn’s multiple comparisons, Monocytes vs. hMDM, **p = 0.0055, Monocytes vs PBMC, p = 0.1302, hMDM vs PBMC, p = 0.7332). As most of the correlations with dopamine receptor expression were in hMDM, these data indicate that the correlations are not driven by higher baseline IL-1β. Together, these data show that dopamine receptor expression varies between types of myeloid cells and changes with maturation. These data also suggest the dopamine receptor levels in hMDM, namely the high D1-like to D2-like ratio, may be important in driving the inflammatory effect of dopamine in this cell type.

### 3.3 Dopamine receptor-dependent effects on IL-1β expression in human microglia

Since we detected differences in dopamine receptor expression between monocytes and MDMs, we speculated that microglia could respond differently from MDMs to dopamine. To answer this question, we used the C06 and C20 human microglial cell lines (Garcia-Mesa, Jay et al. 2017, Alvarez-Carbonell, Ye et al. 2019, Ye, Alvarez-Carbonell et al. 2022, Rheinberger, Costa et al. 2023). These cells express numerous microglial markers, including CD11b, TMEM119, CD68, TGFβR, and P2RY12, maintain migratory capacity and phagocytic activity characteristic of microglia, and expressed P2RY12 and TMEM119 in our culture system (**Supplementary Figure 4**).

Both the C06 and C20 cells expressed mRNA for DRD1, DRD2, and DRD5, but not DRD3 or DRD4. Interestingly, the pattern of dopamine receptor expression was dissimilar between the cell lines. The C20 cells have significantly more DRD1 than the C06 cells (Mann-Whitney test, n = 8, **p = 0.0070, sum of (C06 D1 vs C20 D1) ranks 43, 93, U=7), while the C06 cells have significantly more DRD2 than the C20 cells (Mann-Whitney test, n = 7 - 9, ***p = 0.0002, sum of (C06 D2 vs C20 D2) ranks 91, 45, U=0). There was no difference in baseline DRD5 expression between the lines (Mann-Whitney test, n = 7 - 8, p = 0.6943, sum of (C06 D5 vs C20 D5) ranks 52, 68, U=24). When considering these differences within each line, the C06 cells showed significantly higher DRD2 expression compared to the D1-like receptors, DRD1 and DRD5 (**Figure 3A**, Kruskal-Wallis test, n= 7 - 8, Kruskal-Wallis statistic 8.666, **p = 0.0082; *Post-hoc* with Dunn’s multiple comparisons, D1 vs. D2, * p = 0.0449, D5 vs. D2, * p = 0.0224). Conversely, in C20 cells, there was significantly higher expression of the D1-like receptors compared to DRD2 (**Figure 3B**, Kruskal-Wallis test, n=8-9, Kruskal-Wallis statistic 19.07, ****p < 0.0001; *Post-hoc* with Dunn’s multiple comparisons, DRD1 vs. DRD2, **** p < 0.0001, DRD5 vs. DRD2, * p = 0.0423). When calculating the D1-like to D2-like receptor ratio in both cell lines, we determined that the ratio was 0.29 in C06 cells and 7.7 in C20 cells.

**Figure 3:**
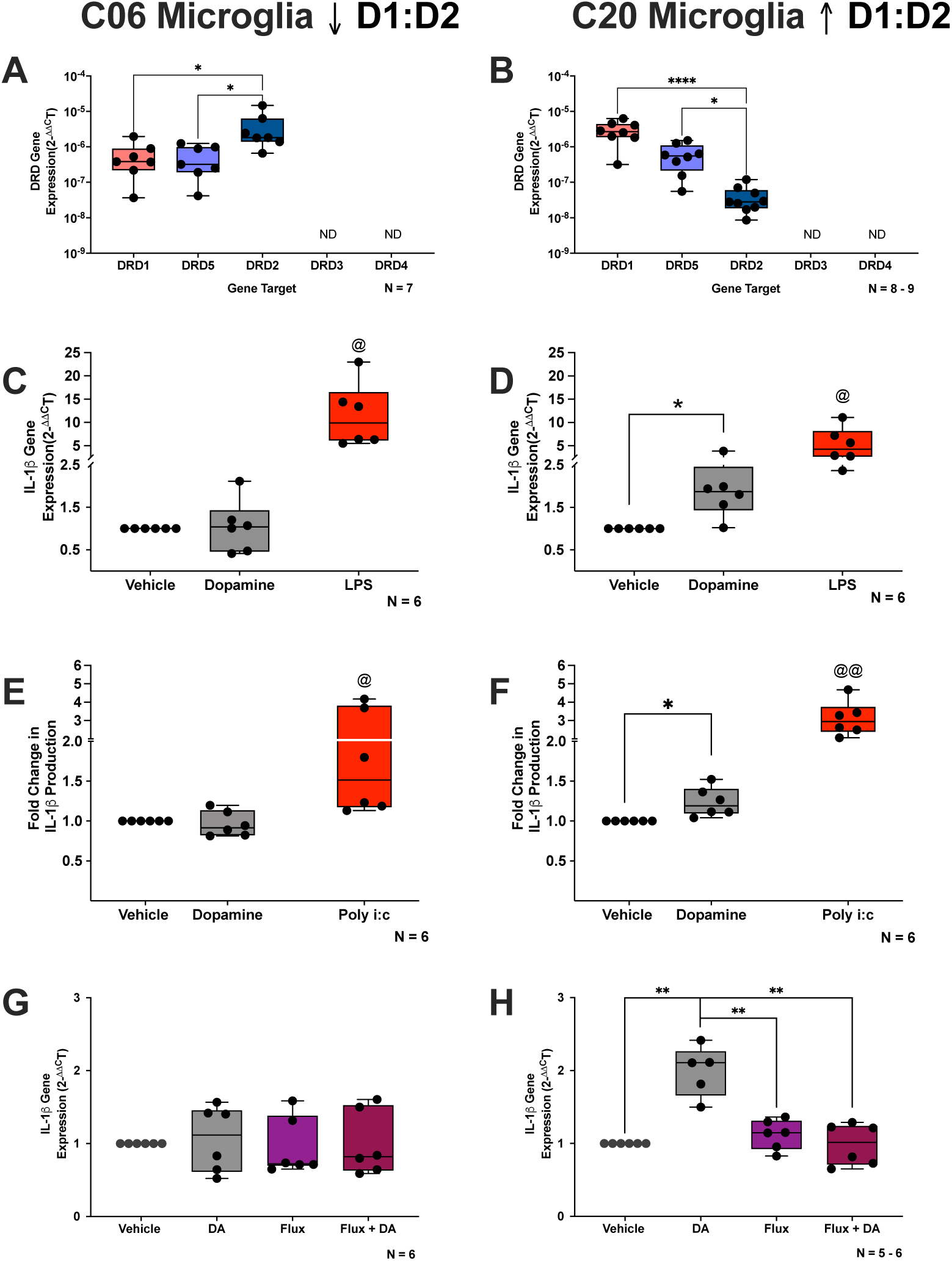
Dopamine receptor-dependent effects on IL-1β expression in human microglia. Dopamine receptor transcripts were measured by qPCR. C06 and C20 microglia express DRD1, DRD2, and DRD5 mRNA. In C06 microglia, there was significantly higher DRD2 expression compared to the D1-like receptors, and in C20 microglia, there was significantly higher expression of the D1-like receptors compared to DRD2. **(C)** C06 and **(D)** C20 microglia were treated with dopamine (10^-6^ M) or LPS (10 ng/mL) as a positive control for 3 hours and examined for IL-1β mRNA. There was no effect in C06 microglia, but in C20 microglia, dopamine significantly increased IL-1β mRNA. **(E)** C06 and **(F)** C20 microglia were treated with dopamine (10^-6^ M) or poly I:C (10 ug/mL) as a positive control for 4 hours, then lysates were collected and examined for IL-1β production by AlphaLISA. There was no effect in C06 microglia. However, in C20 microglia dopamine significantly increased IL-1β production. **(G)** C06 and **(H)** C20 microglia were pretreated with flupentixol (Flux, 10-6 M) or vehicle for 30 min in the presence of absence of 3 hour dopamine (DA, 10^-6^ M) treatment and examined for IL-1β mRNA. There was no effect of treatment on C06 microglia, but in C20 microglia Flux significantly blocked the DA-mediated increase in IL-1β. Significance was determined by paired t-tests, Wilcoxon tests, Kruskal-Wallis tests, post-hoc with Dunn’s multiple comparisons, and a two-way mixed effects model and post-hoc with Tukey’s multiple comparisons, *p < 0.05, **p < 0.01, ***p < 0.001, and *****p < 0.0001.

We then took advantage of the differential D1-like to D2-like receptor ratios in the two lines to enable a cleaner comparison of the specific impact of dopamine receptor subtypes on dopamine-mediated IL-1β. The C06 and C20 cells were treated with dopamine (10^-6^ M) or LPS (10 ng/mL, positive control) for 3 hours and examined for IL-1β mRNA expression. In the C06 cells, which have a low D1-like to D2 ratio, dopamine did not increase IL-1β mRNA (**Figure 3C**, Vehicle vs Dopamine, n = 6, paired t-test, p = 0.2614, t = 1.266, df = 5). However, in the C20 cells, with a high D1-like to D2 ratio, dopamine significantly increased IL-1β mRNA (**Figure 3D**, Vehicle vs Dopamine, n = 6, Wilcoxon test, *p = 0.0312, sum of (+,-) ranks 21, 0). LPS significantly increased IL-1β expression in both cell lines (**Figure 3C-D**, C06 IL-1β Vehicle vs LPS, n = 6, paired t-test, ^@^p = 0.0114, t = 3.901, df = 5; C20 IL-1β Vehicle vs LPS, n = 6, Wilcoxon test, ^@^p = 0.0312, sum of (+,-) ranks 21, 0), demonstrating that C06 cells still have the capacity to increase IL-1β mRNA similar to C20 cells.

To verify that the effect of dopamine was impacting IL-1β production and not only transcription, the C06 and C20 microglia were treated with dopamine (10^-6^ M) for 4 hours, then lysates were collected and examined for IL-1β production. As with mRNA, in the C06 cells (low D1-like to D2 ratio) dopamine treatment did not significantly increase IL-1β production (**Figure 3E**, Vehicle vs Dopamine, n = 6, Wilcoxon test, p = 0.6875, sum of (+,-) ranks 8, -13). However, in the C20 cells (high D1-like to D2 ratio), dopamine did significantly increase IL-1β production, again in line with the effect on mRNA (**Figure 3F**, Vehicle vs Dopamine, n = 6, paired t-test, *p = 0.0384, t = 2.790, df = 5). In these experiments, Poly I:C treatment was used as a positive control, and it significantly increased expression in both cell lines (**Figure 3E-F**, C06 IL-1β Vehicle vs Poly I:C, n = 6, Wilcoxon test, ^@^p = 0.0312, sum of (+,-) ranks 21, 0; C20 IL-1β Vehicle vs Poly I:C, n = 6, paired t-test, ^@@^p = 0.0060, t = 4.564, df = 5). Thus, dopamine-mediated increases in IL-1β production in C20 cells mirrored the change in IL-1β mRNA levels (increase in C20 IL-1β Production vs increase C20 IL-1β mRNA, n=6, unpaired t-test with Welch’s correction, p=0.1005, t=1.981, df=5.361). This indicates that dopamine impacts IL-1β gene and protein expression similarly in these cells and therefore we used mRNA expression as a readout in the remainder of these studies.

Although the primary mechanism by which dopamine mediates its effects is activation of its cognate receptors, dopamine has also been shown to act on immune cells via activation of adrenergic receptors or through formation of reactive oxygen species (Channer, Matt et al. 2022). To confirm that the inflammatory effects of dopamine are dependent on dopamine receptor activation, C06 and C20 cells were pretreated with the pan dopamine receptor antagonist flupentixol (Flux, 10^-6^ M) for 30 minutes followed by 3-hour dopamine (DA, 10^-6^ M) treatment. We detected no effect of either dopamine or flupentixol on IL-1β mRNA expression in C06 microglia (**Figure 3G**, rm 2-way ANOVA, Flupentixol, p = 0.4440 F (1,5) = 0.6902, Dopamine, p= 0.3358 F (1,5) = 1.133, Interaction, p = 0.6157 F (1,5) = 0.2860). In contrast, C20 microglia showed a significant, main effect of both flupentixol and dopamine, as well as an interaction between flupentixol and dopamine, in that flupentixol eliminated the dopamine-mediated increase in IL-1β. (**Figure 3H**, rm 2-way mixed effects model and *post-hoc* with Tukey’s multiple comparisons, Flupentixol, **p = 0.0013 F (1,5) = 42.08, Dopamine, *p= 0.0203 F (1,5) = 11.23, Interaction, ***p = 0.0008 F (1,4) = 81.01, No Flux:Vehicle vs No Flux:Dopamine, **p = 0.0035, No Flux:Dopamine vs Flux:Vehicle, **p = 0.0031, No Flux:Dopamine vs Flux:Dopamine, **p = 0.0092). These data confirm that the effect of dopamine on IL-1β in C06 and C20 cells is mediated via dopamine receptors.

To determine how the D1-like to D2-like dopamine receptor ratio mediates regulation of IL-1β, C20 cells were transduced with a TetR lentivirus followed by a DRD2 lentivirus, creating a cell line in which DRD2 was overexpressed in response to tetracycline (C20D2). To validate the model, the C20D2 cells were exposed to tetracycline and examined for dopamine receptor mRNA expression and compared to wild type (WT) C20 cells. In response to tetracycline, the C20D2 cells expressed significantly more DRD2 than WT C20 microglia (**Supplementary Figure 5A**, Kruskal-Wallis test, n = 7 - 9, Kruskal-Wallis statistic 18.49, ****p < 0.0001; *Post-hoc* with Dunn’s multiple comparisons WT DRD2 vs. C20D2 +Tet DRD2, ****p < 0.0001), while also expressing significantly less DRD1 than WT C20 microglia (**Supplementary Figure 5B**, one-way ANOVA, n = 7 - 8, ***p = 0.0005, F (2, 19) = 7.106; *Post-hoc* with Tukey’s multiple comparisons, WT DRD1 vs. C20D2 +Tet DRD1, **p = 0.0012). Even in the absence of tetracycline, the C20D2 cells showed significantly less DRD1 (WT DRD1 vs. C20D2 -Tet DRD1, **p = 0.0017) and trended toward less DRD2 (WT DRD2 vs. C20D2 -Tet DRD2, p = 0.0765). No significant differences were observed in expression of DRD5 (**Supplementary Figure 5C**, one-way ANOVA, n = 7 - 8, p = 0.1163, F (2, 19) = 1.380).

As anticipated the overexpression of DRD2 altered the D1-like to D2-like receptor ratio relative to the WT C20 cells. The WT C20 had a ratio of 7.7, while the non-tetracycline exposed C20D2 cells had a ratio of 3.2 and the tetracycline treated C20D2 cells had a ratio of 0.08. This is very close to that of the WT C06 cells, which had a D1-like to D2-like receptor ratio of 0.29 and indicates a successful shift of the dopamine receptor ratio to resemble the non-dopamine responsive WT C06 cells. Next, we treated the WT C20, the non-tetracycline treated C20D2 and the tetracycline treated C20D2 cells with dopamine (10^-6^ M) for 3 hours and examined IL-1β mRNA. The WT C20 cells (D1-like to D2-like 7.7) showed a 2-fold increase in IL-1β, the non-tetracycline treated C20D2 (D1-like to D2-like 3.2) showed a 1.73 fold increase in IL-1β, and the tetracycline treated C20D2 (D1-like to D2-like 0.08) showed a 1.41 fold increase in IL-1β. Although the difference was not significant (**Supplementary Figure 7D**, one-way ANOVA, n = 6 - 7, p = 0.3542, F (2, 17) = 0.6817), the level of response to dopamine directly correlated with the D1-like to D2-like receptor ratio, supporting our hypothesis that the ratio of D1-like receptors to D2-like receptors is important for dopamine’s effects on inflammation.

### 3.4 Differential gene expression, pathway analysis and upstream regulators in dopamine-treated human microglia

To identify genes and pathways that are activated in dopamine treated C20 cells, we performed RNA-seq. We used a cutoff value of fold change > |1.1| and FDR < 0.05 for inclusion in our analyses and there were a total of 1604 genes that were differentially expressed between the vehicle and dopamine treated C20 cells (**Figure 4A**). Of these 776 genes were increased and 828 genes were decreased (**Supplementary Table 2**). Importantly, these analyses confirmed our mRNA analyses, showing that IL-1β was increased 2.23-fold in dopamine-treated C20 cells, and identified increases in genes that have previously been found to be upregulated [KYNU (4.76-fold), RASD1 (3.51-fold), and PDE4B (2.33-fold)] by dopamine and dopamine receptor agonists in the human U1 monocytic cell line (Basova, Lindsey et al. 2023) (**Figure 4B**, data expressed as transcripts per million (TPM)). Moreover, these data showed that dopamine impacted numerous regulators of inflammation, dopaminergic signaling, and other processes in this microglial cell line.

**Figure 4:**
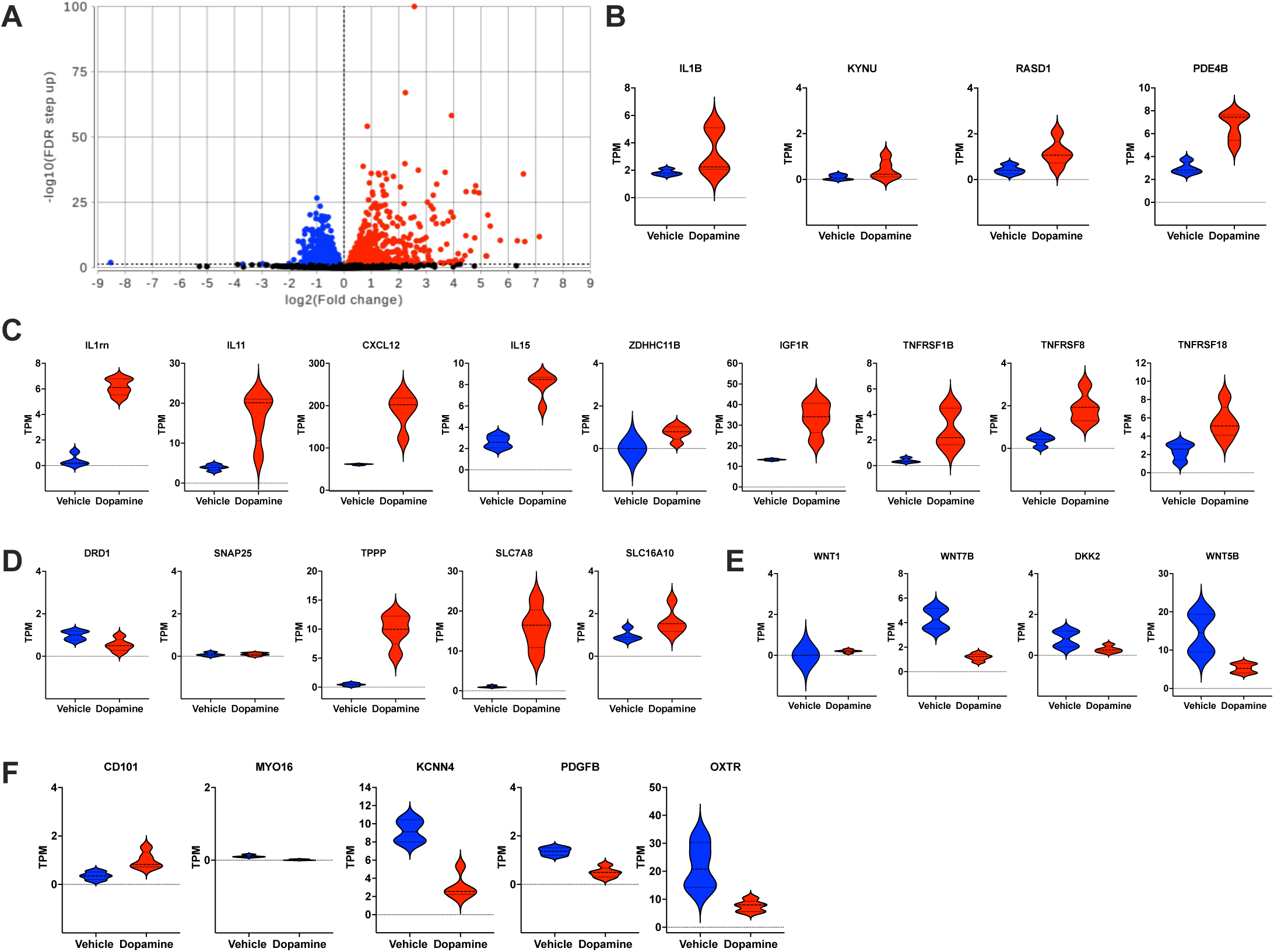
Differential gene expression of dopamine-treated human microglia. **(A)** Dopamine induced both up-regulated (776) and down-regulated (828) genes (fold change > |1.1| and FDR < 0.05) in C20 microglia. Violin plots showing the relative expression of genes in the two different conditions (vehicle or dopamine), including **(B)** genes previously found to be upregulated by dopamine in myeloid cells, **(C)** genes associated with modulation inf inflammation, **(D)** genes associated with regulation of dopaminergic signaling, **(E)** genes associated with Wnt signaling, and **(F)** genes associated with microglia identity and activation.

Some of the genes associated with the modulation of inflammation include; the cytokines and chemokines IL1RN (21.96 fold), IL11 (4.47-fold), CXCL12 (2.83-fold) and IL15 (2.66-fold); ZDHHC11B (52.17-fold), a positive modulator in NF-κB signaling (Liu, Sun et al. 2021); IGF1R (2.59-fold), which can alter microglial polarization via TLR4/NF-κB pathway (Sun, Wu et al. 2020) and several tumor necrosis factor (TNF) receptors, TNFRSF1B (TNFR2, CD120b, 7.41-fold) which binds to TNF-α and is neuroprotective (Pillai, Bebek et al. 2021), TNFRSF8 (CD30, 5.38-fold) which binds to CD153 and can activate NF-κB signaling (Wright and Duckett 2009), and TNFRSF18 (GITR, CD357, 2.52-fold) which binds to GITR ligand and is involved in the pathogenesis of autoimmune disease (Tian, Zhang et al. 2020) (**Figure 4C**). Examples of the genes associated with the regulation of dopaminergic signaling include; DRD1 itself (-2.12-fold), which may be downregulated in response to dopamine-mediated activation, SNAP25 (6.62-fold), which is important for neurotransmitter release and has been previously identified inside microglial processes following synaptic contacts (Paolicelli, Bolasco et al. 2011); TPPP (28.02-fold), which regulates plasma membrane presentation of the dopamine transporter and enhances cellular sensitivity to dopamine toxicity; and the transporters SLC7A8 (15.18-fold), an Na+-independent transporter of neutral amino acids that transports L-DOPA and SLC16A10 (2.16-fold), which mediates Na-independent transport of tryptophan, tyrosine, phenylalanine, and L-DOPA (**Figure 4D**). There were also dopamine-mediated changes in genes associated with Wnt signaling [WNT1 (12.54-fold), WNT7B (-3.19-fold), DKK2 (-2.87-fold), and WNT5B (-2.85-fold)], which is important for normal dopaminergic functioning as well as inflammation and neurodegeneration during dopaminergic dysfunction (Marchetti and Pluchino 2013, L’Episcopo, Tirolo et al. 2018) (**Figure 4E**). Last, a number of genes were altered that are associated with microglia identity and activation [CD101 (2.76-fold), MYO16 (-7.86-fold), KCNN4 (-2.80-fold), PDGFB (-3.04-fold), and OXTR (-2.82-fold)] (Kaushal, Koeberle et al. 2007, Yuan, Liu et al. 2016, Felsky, Roostaei et al. 2019, Bi, Wang et al. 2022, Selles, Fortuna et al. 2023) (**Figure 4F**).

To evaluate how these alterations might impact production of IL-1β and other inflammatory pathways, we mapped the changes in gene expression onto known patterns in canonical pathways using Ingenuity Pathway Analysis (IPA) (**Supplementary Table 3**). The pathways with the greatest Z-score that showed an increase in dopamine treated cells relative to vehicle were IL-10 Signaling, Protein Kinase A Signaling, PTEN Signaling, PPARα/RXRα Activation, Activin Inhibin Signaling Pathway, all of which mediate inflammation and most of which involve IL-1β. The pathways with the greatest Z-score that showed a decrease in dopamine treated cells relative to vehicle included Neuroinflammation Signaling Pathway, IL-8 Signaling, iNOS Signaling, PI3K/AKT Signaling, HIF1α Signaling, Role of PKR in Interferon Induction and Antiviral Response, Chemokine Signaling, NF-κB Activation by Viruses, and IL-1 Signaling.

We also used IPA upstream regulator analysis to identify the potential upstream regulators for these pathways based on expected effects between transcriptional regulators and the targets. This predicts relevant transcriptional regulators by examining how many targets are present in the data and compares the direction of changes in these targets to what is defined the literature. If the direction of changes is mostly consistent with a particular activation state, a prediction is made with two statistical measures: the p-value of overlap and Z-score (Krämer, Green et al. 2014), where a positive Z-score indicates activation and a negative Z-score indicates inhibition. This process identified several regulators which are listed in **Supplementary Table 4**. These include but are not limited to upstream regulators associated with neuronal activity and neurotransmission [NEUROD1 (Z-score 2.373), norepinephrine (Z-score 2.946), FOS (Z-score -2.026), FOSB (Z-score -2.2), and SLC16A2 (Z-score -2.074)], microglia-enriched transcription factors [SAL4 (Z-score 3.889), RARG (Z-score 2.186) and TEAD4 (Z-score -2.005)] and inflammatory signaling [CD247 (Z-score 2.236), IL9 (Z-score 2.229), TLR9 (Z-score 2.155), AMPK (Z-score 2.08), FHL2 (Z-score -2.804), IL3 (Z-score -2.428), CD40 (Z-score -2.382), HMGXB4 (Z-score -2.236), PTPN2 (Z-score -2.219), CD40LG (Z-score -2.21), CCL5 (Z-score -2.179), LTA (TNFβ) (Z-score -2.138), TIMP1 (Z-score -2.111), and NFKB1 (Z-score -2.015)]. Some notable effects were the activation of norepinephrine, the microglia-enriched transcription factors RARG and TEAD4 (Ayata, Badimon et al. 2018) and the inhibition of the negative regulator of Wnt signaling, HMGXB4 (He, Dong et al. 2021), as well as of NFKB1 and the negative regulators of NF-kB and inflammasome activation, FHL2 and PTPN2 (Ha Thi, Choi et al. 2016, Dahan, Levillayer et al. 2017). These experiments demonstrated a significant interaction between dopamine and the regulation of inflammation in human microglia and suggest further research in this area is needed to better define these effects.

### 3.5 Dopamine-induced IL-1β Exacerbated with HIV

Dopamine mediated increases in IL-1β would be most relevant in the context of chronic inflammatory disease with highly comorbid substance use disorders (SUDs). One of the diseases that carries this type of SUD burden is infection with Human Immunodeficiency Virus (HIV), in which comorbid SUD is the among the most common neuropsychiatric comorbidities (Nedelcovych, Manning et al. 2017, Cook, Burke-Miller et al. 2018, Heslin, Jewell et al. 2023). Our prior data in human macrophages and microglia also show that exposure to dopamine concentrations evoked by the use of psychostimulants (10^-6^ M) significantly increase HIV viral replication in hMDM, C06/C20 microglia, and iPSC-derived human microglia (Gaskill, Calderon et al. 2009, Gaskill, Yano et al. 2014, Nickoloff-Bybel, Mackie et al. 2019, Matt, Nickoloff-Bybel et al. 2021) (representative images in **Figure 5A** and **6C)**, and that this is specifically driven by activation of dopamine receptors (Gaskill, Yano et al. 2014, Nickoloff-Bybel, Mackie et al. 2019). Importantly, dopamine can also increase the production of inflammatory cytokines and chemokines in virally suppressed, chronically infected HIV patients (Nolan, Muir et al. 2018), demonstrating a strong connection between HIV infection, inflammation, and dopaminergic dysfunction, with a specific focus in myeloid populations.

**Figure 5:**
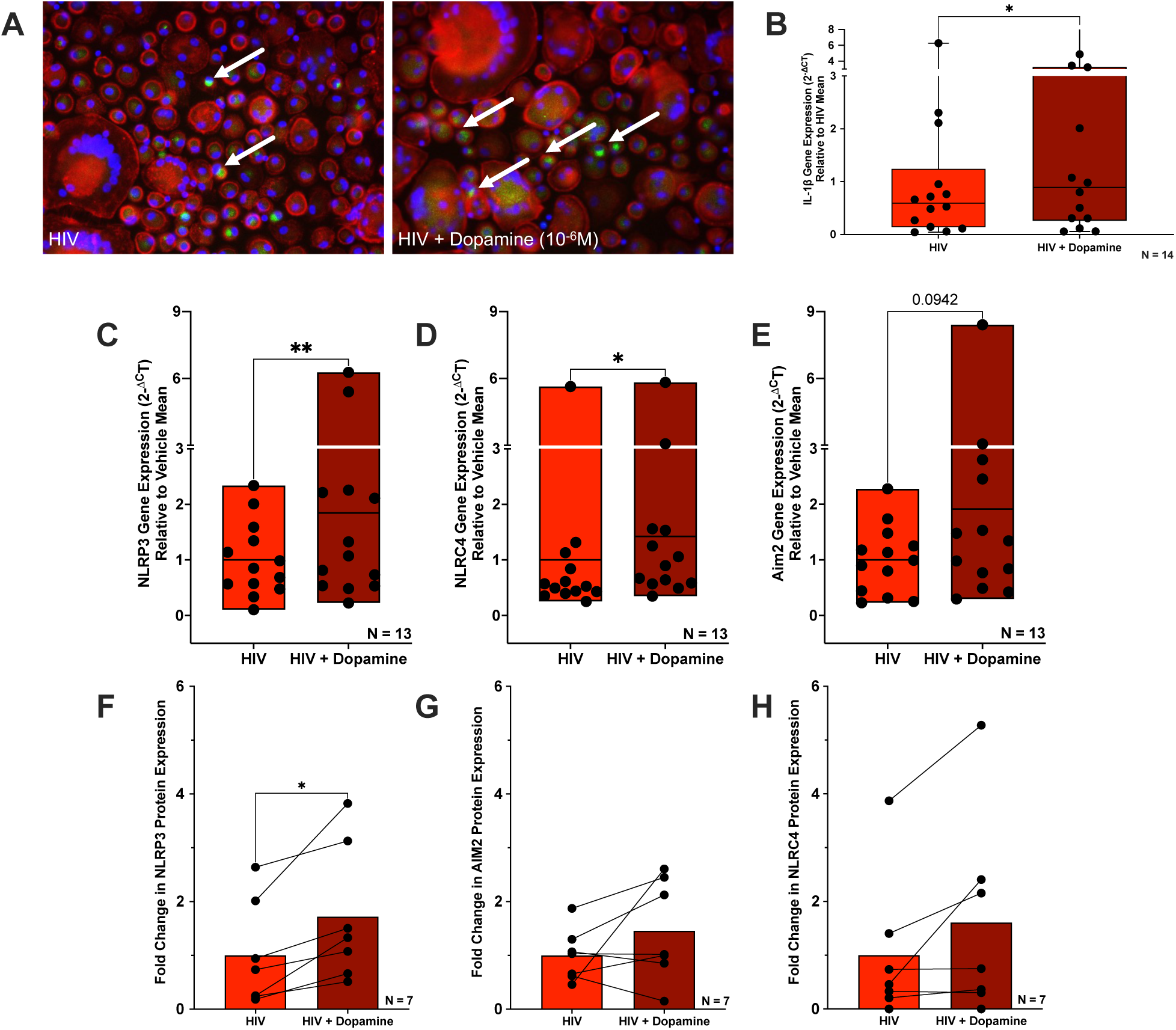
Dopamine increases IL-1β and inflammasome expression in HIV-infected primary macrophages. **(A)** Representative immunofluorescent images at 20X objective (DAPI, blue; Phalloidin, red; p24, green) show human monocyte-derived macrophages (hMDM) infected with 0.5 ng/ml HIV_ADA_ and treated with dopamine (10^-6^ M) for 7 days. Higher levels of cell fusion and giant cell formation as well as p24 expression (white arrows) are seen in infected dopamine-treated cultures relative to cultures only infected with HIV. hMDM were also infected with 0.5 ng/ml HIV_ADA_ for 7 days and then treated with dopamine (10^-6^ M) for 3 hours and examined for **(B)** IL-1β, **(C)** NLRP3, **(D)** NLRC4, and **(E)** AIM2 expression by qPCR. There was a significant increase in IL-1β, NLRP3, and NLRC4 and a trending increase in AIM2 expression with HIV + Dopamine relative to HIV alone. With these same samples we simultaneously extracted protein and examined **(F)** NLRP3, **(G)** NLRC4, and **(H)** AIM2 expression by Western Blot. There was only a significant increase in NLRP3 expression with HIV + Dopamine relative to HIV alone. Significance was determined by paired t-tests and Wilcoxon tests, *p < 0.05 and **p < 0.01.

**Figure 6:**
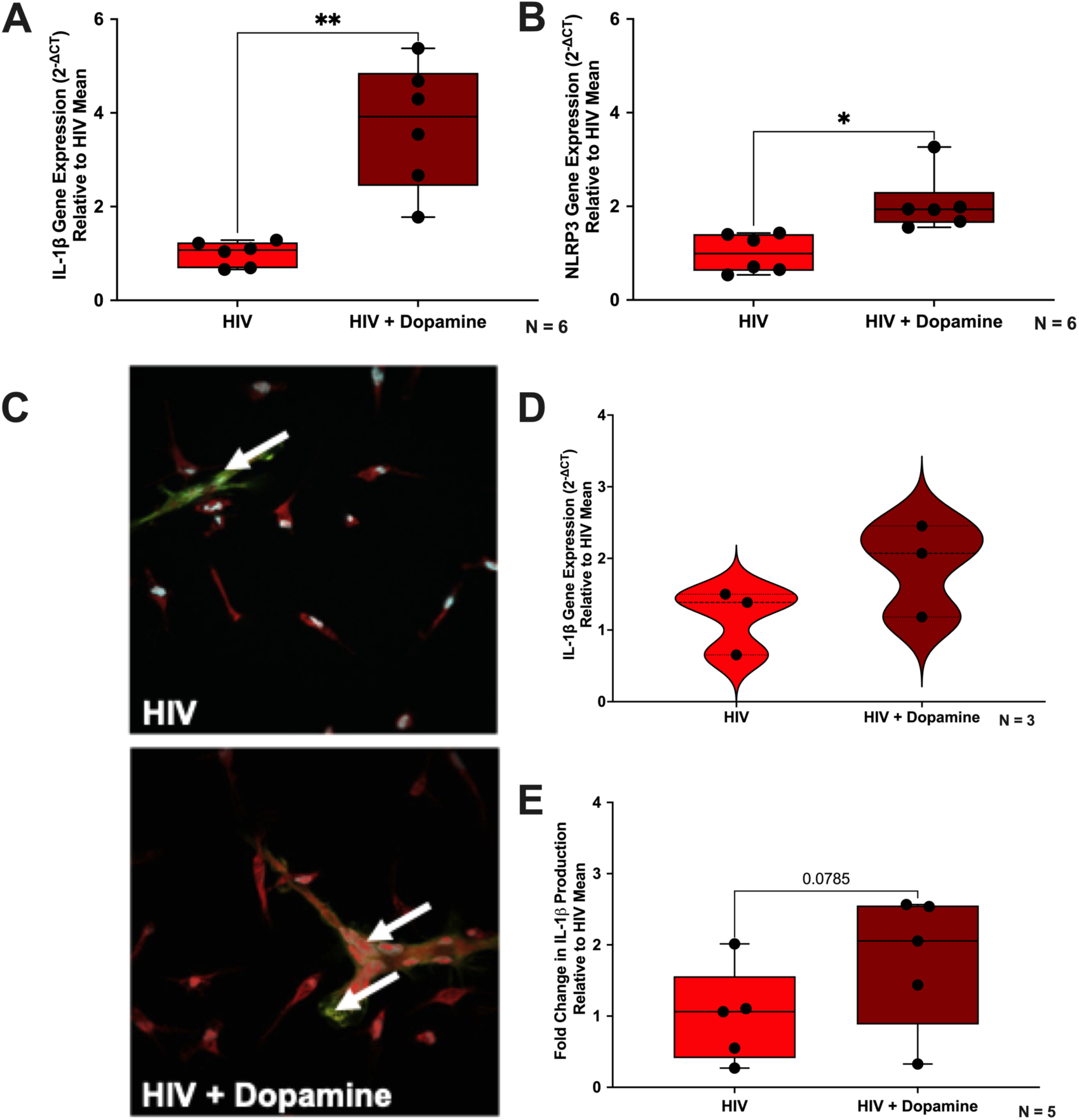
Dopamine increases IL-1β and NLRP3 in HIV-infected microglia. C20 microglia were infected with 2.5 ng/ml HIV_ADA_ for 48 hours, then treated with dopamine (10^-6^ M) for 3 hours and examined for **(A)** IL-1β and **(B)** NLRP3 mRNA expression by qPCR. For both IL-1β and NLRP3 there was a significant increase with HIV + Dopamine relative to HIV alone. **(C)** iMicroglia were infected with 1 ng/mL of HIV_ADA_ for 7 days with or without dopamine (10^-6^ M) and representative immunofluorescent images at 60X objective (DAPI, blue; CellMask, red; p24, green) show higher levels of cell fusion and giant cell formation as well as p24 expression (white arrows) in infected dopamine-treated cultures relative to cultures only infected with HIV. iMicroglia were also infected with 1 ng/mL of HIV_ADA_ for 7 days and then treated with dopamine (10^-6^M) for either 3 hours to examine IL-1β mRNA expression with qPCR or 4 hours to examine IL-1β production by AlphaLISA. **(D)** Analysis of IL-1β mRNA levels show dopamine increases IL-1β in HIV-infected cells. **(E)** Analysis of lysate IL-1β levels by AlphaLISA show there is a trend that dopamine increases IL-1β in HIV-infected cells. Significance was determined by paired t-tests and Wilcoxon tests, *p < 0.05 and **p < 0.01.

To determine how HIV infection impacts the dopamine-mediated increase in IL-1β in myeloid cells, hMDM were infected for 7 days with 0.5 ng/mL HIV_ADA_, treated with dopamine (10^-6^ M) for 3 hours and examined for IL-1β mRNA expression. Treatment with dopamine significantly increased IL-1β expression in HIV-infected hMDM relative to infected cells treated with vehicle (**Figure 5B**, Wilcoxon test, n = 14, * p = 0.0107, sum of (+,-) ranks 92, -13). The magnitude of this effect, an almost 70% increase, was nearly 2-fold greater than what we have previously observed in uninfected hMDM (Nolan, Reeb et al. 2020). Expression of NLRP3 paralleled the trend seen with IL-1β, with the dopamine-treated, HIV-infected cells showing significantly higher NLRP3 mRNA expression compared to vehicle-treated, HIV-infected cells (**Figure 5C**, Wilcoxon test, n = 13, ** p = 0.0061, sum of (+,-) ranks 83, -8). As some data indicate that activation of inflammasomes by different HIV proteins can act discretely on IL-1β and IL-18 secretion (Triantafilou, Ward et al. 2021), we also examined the effects of dopamine on 2 other inflammasomes, NLRC4 and AIM2. Dopamine-treated, HIV-infected cells showed significantly higher NLRC4 mRNA expression (**Figure 5D**, Wilcoxon test, n = 13, * p = 0.0327, sum of (+,-) ranks 76, -15) and a trend towards higher AIM2 mRNA expression (**Figure 5E**, Wilcoxon test, n = 13, p = 0.0942, sum of (+,-) ranks 49, 13) compared to vehicle-treated, HIV-infected cells. These results are closely mirrored in the dopamine effects on NLRP3, NLRC4, and AIM2 protein expression in these cells. The dopamine-treated, HIV-infected cells had higher NLRP3, NLRC4, and AIM2 protein expression compared to vehicle-treated, HIV-infected cells, although this only reached significance for NLRP3 (**Figure 5F**, NLRP3 paired t-test, n = 7, *p = 0.0133, t = 3.471, df = 6; **Figure 5G**, NLRC4 Wilcoxon test, n = 7, p = 0.0938, sum of (+,-) ranks 19, -2, **Figure 5H**, AIM2 paired t-test, n = 7, p = 0.2143, t = 1.389, df = 6). Representative Western blots are shown in **Supplementary Figure 6**.

We then performed similar experiments using C06 and C20 microglia to better define the precise effect of HIV infection in CNS myeloid populations. Both C06 and C20 microglia were infected with 2.5 ng/mL HIV_ADA_ for 48 hours, then treated with dopamine (10^-6^ M) for 3 hours and examined for IL-1β mRNA expression. As in the hMDM, dopamine treatment of HIV-infected C20 cells significantly increased the level of IL-1β compared to HIV infection alone (**Figure 6A**, HIV vs HIV + Dopamine, n = 6, paired t-test, **p = 0.0048, t = 4.811, df = 5). This was also nearly a 2-fold greater increase relative to the uninfected dopamine-mediated increase seen in C20 cells in **Figure 3**. As anticipated based on the data in **Figure 3**, dopamine did not show any effect on IL-1β on HIV infected C06 microglia (HIV vs HIV + Dopamine, n = 6, paired t-test, p = 0.1954, t = 1.494, df = 5), so the remainder of these experiments focused only on the C20 microglia. Like hMDM, expression of NLRP3 paralleled the trend seen with IL-1β, with the dopamine-treated, HIV-infected cells showing significantly higher NLRP3 mRNA expression compared to vehicle-treated, HIV-infected cells (**Figure 6B**, HIV vs HIV + Dopamine, n = 6, Wilcoxon test, * p = 0.0312, sum of (+,-) ranks 21, 0). We also examined other inflammasome associated genes in HIV-infected C20 cells, and while dopamine did slightly increase the amount of CASP1 or PYCARD mRNA in these cells, these effects were not significant (**Supplementary Figure 7C**, HIV vs HIV + Dopamine, n = 6, Wilcoxon test, p = 0.3125, sum of (+,-) ranks 16, -5; **Supplementary Figure 7D**, HIV vs HIV + Dopamine, n = 6, paired t-test, p = 0.1890, t = 1.520, df = 5). Examining the scaffolding proteins for other types of inflammasomes, we found that there was a significant dopamine-mediated decrease in NLRP1 (**Supplementary Figure 7E**, HIV vs HIV + Dopamine, n = 6, paired t-test, **p = 0.0035, t = 5.194, df = 5), and no significant change in NLRC5 (**Supplementary Figure 7F**, HIV vs HIV + Dopamine, n = 6, paired t-test, p = 0.4395, t = 0.8394, df = 5). There was no detectable expression of AIM2 and NLRC4 mRNA in the C20 microglial cell line.

Surprisingly, the effect of dopamine in microglia impact other IL-1 family cytokines, as HIV-infected C20 cells showed a trend toward dopamine decreasing IL-18 mRNA expression (**Supplementary Figure 7A**, HIV vs HIV + Dopamine, n = 6, paired t-test, p = 0.0861, t = 2.133, df = 5). We also examined IL-1 receptor antagonist (IL1rn), a naturally occurring inhibitor of IL-1β which has also been shown to be upregulated by HIV (Zavala, Rimaniol et al. 1995, Kreuzer, Dayer et al. 1997, Nouhin, Pean et al. 2017). and found a 25-fold increase in this gene (**Supplementary Figure 7B**, HIV vs HIV + Dopamine, n = 6, paired t-test, ***p = 0.0003, t = 8.922, df = 5), corroborating the 22-fold increase in IL1rn seen in our RNA-seq data.

To verify these findings in a model that more closely match human microglia, we repeated our experiments using iPSC-derived microglia (iMicroglia). These iMicroglia have very similar gene expression to primary human microglia (Ryan, Gonzalez et al. 2020) and express the microglial markers TMEM119, P2RY12, and Iba1 in our culture system (**Supplementary Figure 8**). We have previously shown that iMicroglia express the D1-like dopamine receptors, DRD1 and DRD5, but not D2-like receptors, and that dopamine significantly increases HIV infection in these cells (Matt, Nickoloff-Bybel et al. 2021). In these experiments, iMicroglia were mock-infected or infected with 1 ng/mL of HIV_ADA_ for 7 days and then treated with vehicle (H_2_O) or dopamine (10^-6^M), treated for either 3 hours to examine IL-1β mRNA expression or 4 hours to examine IL-1β production. Dopamine increased IL-1β mRNA by 2.65-fold (**Figure 6D**, HIV vs HIV + Dopamine, n = 3, paired t-test, p = 0.3940, t = 1.077, df = 2). Analysis of intracellular IL-1β production showed dopamine increased IL-1β in HIV-infected cells by 2.06-fold (**Figure 6E**, HIV vs HIV + Dopamine, n = 5, paired t-test, p = 0.0785, t = 2.350, df = 4), and found that the increase with dopamine is greater in HIV-infected compared to uninfected cells but is not significant (data not shown, Mock-infected vs HIV-infected dopamine-mediated fold change in IL-1β, n = 5, paired t-test, p = 0.3721, t = 1.004, df = 4). These data, and the similarity of these effects across all three myeloid models, indicate that dopamine can enhance IL-1β expression in HIV-infected human macrophages and microglia, and suggest that increased levels of dopamine due to substance misuse or other therapeutics could exacerbate inflammation associated with neuroHIV.

## 4. Discussion

Here we demonstrate that increased production of IL-1β by dopamine depends on the distinct dopamine receptor expression on several types of human myeloid cells. hMDM expressing DRD1 showed a significant, dopamine-mediated increase in IL-1β that was not present in hMDM without DRD1, suggesting this receptor is necessary for the increase. A similar effect was seen for expression of DRD4. And while dopamine-mediated increases in IL-1β were seen in hMDM both expressing and not expressing DRD2 and DRD3, cells lacking these receptors had a greater magnitude of dopamine-mediated increases in IL-1β, suggesting that they might be more associated with mitigating inflammation. A similar effect was observed in human microglial cell lines, which showed that dopamine only increased IL-1β gene and protein expression in microglia with a high ratio of D1-like receptors relative to DRD2. Antagonizing all dopamine receptors on microglia by pre-treatment with flupentixol (pan-dopamine receptor antagonist) confirmed the specificity of these effects to dopamine receptors. Unlike in hMDM, the microglial cell lines showed these effects without the presence of DRD3 and DRD4, suggesting that DRD2 may be the primary driver of anti-inflammatory effects in these cells. Notably, our results suggest that DRD1, a low-affinity receptor, drives pro-inflammatory activity in hMDM while DRD2, a high-affinity receptor, drives anti-inflammatory activity. This is supported by studies showing that dopamine preferentially binds to D2-like receptors at low concentrations and D1-like receptors at high concentrations (Richfield, Penney et al. 1989, Martel and Gatti McArthur 2020). However, the connection between dopamine receptor expression and IL-1β activity is likely to be more complicated than this. Dopamine receptor affinity for dopamine does not break down strictly by sub-type, as the affinity of DRD5 is higher than that of DRD1 and DRD2 (Gaskill, Calderon et al. 2013). Further, in hMDM, DRD4 and DRD3, which both have high affinity for dopamine, had opposing effects on dopamine-mediated changes in IL-1β. Some of these differences could be associated with receptor reserve, as this plays a role in the immune cell response to neurotransmitters including dopamine and has been established in T cells (Meller, Bohmaker et al. 1987, McNeil and Evavold 2003, Guieu, Brignole et al. 2021). If the expression of dopamine receptor subtypes varies between donors, and amounts of specific dopamine receptors affect IL-1β production, there could be a significant number of interactions associated with each dopamine-mediated effect.

In this study, we suggest that some of the differences in dopaminergic immunoregulation may also be due to changes in dopamine receptor ratios with maturation, as we show distinct dopamine receptor ratios in human macrophages, monocytes and PBMCs. Our data indicate that hMDM have a significantly greater D1-like / D2-like dopamine receptor ratio than either monocytes or PBMC, and the correlations between dopamine receptor expression and IL-1β were strongest in hMDM. This was true despite the finding that monocytes expressed significantly greater baseline IL-1β mRNA than did hMDM, differing from a prior study showing the greatest cytokine expression in human PBMC (Awad, Assrawi et al. 2017). This may reflect the fact that monocytes more rapidly release IL-1β, as they do not need a second signal for inflammasome activation (Netea, Nold-Petry et al. 2009). Our data on dopamine receptor expression also correlate well with a prior study assessing dopamine receptor mRNA expression in monocytes during maturation (Coley, Calderon et al. 2015). Coley and colleagues showed that the mRNA and protein expression of both DRD1 and DRD5 increase as monocytes mature, while DRD4 mRNA and protein decreased with maturation. These data may indicate that the immune cell dopaminergic profile differs between circulating myeloid cells and tissue myeloid cells. Further, they suggest that the high D1-like / D2-like ratio and effects of dopamine on IL-1β would be more prominent in tissues where dopamine levels are higher, as compared to the relatively low dopamine levels found in circulation (Matt and Gaskill 2019). One tissue in which this is clearly the case is the CNS, where the higher levels of dopamine could impact inflammasome activity and IL-1β production in microglia and other CNS-associated macrophages. These effects could also be compounded in individuals with substance use disorders, as chronic use of stimulants, opioids, nicotine, and alcohol not only alter dopamine release, but can selectively decrease DRD2 availability in the brain across multiple species (Volkow, Chang et al. 2001, Nader, Morgan et al. 2006, Fehr, Yakushev et al. 2008). This could bias signaling toward D1-like receptors to increase neuroinflammation and potentially promote inflammation-associated addiction behaviors. Prior studies have shown changes in dopamine receptor expression in peripheral blood cells in individuals with SUD (Czermak, Lehofer et al. 2004, Biermann, Bonsch et al. 2007, Goodarzi, Vousooghi et al. 2009). Whether these changes in peripheral immune cells are related to changes in CNS immune function or the development of SUD remains to be further explored, as this could suggest a better understanding of the immune system’s role in the susceptibility to SUD.

To examine these connections in more detail, we performed RNA-sequencing on C20 microglia-like cells. Several genes associated with inflammatory and neurotransmitter signaling were differentially expressed between vehicle and dopamine-treated cells. Importantly, the effects of dopamine on IL-1β and associated genes (i.e. IL-1rn) were retained in this dataset. Increases in KYNU, RASD1, and PDE4B matched previous data in dopamine-treated myeloid cells (Basova, Lindsey et al. 2023), and we further found notable increases in cytokines/chemokines, TNF receptors, modulators of NF-κB signaling, and amino acid transporters, as well as decreases in Wnt signaling and microglial activation markers. Notably, many altered genes were associated with NF-κB signaling, and an NF-κB signature has been shown to be one factor differentiating microglia from other tissue-macrophages (Batista, Still et al. 2020). This suggests that differences in NF-κB gene expression could be one reason for functional differences between microglia and macrophages regarding their inflammatory response to dopamine. Evaluation of upstream targets using IPA analysis indicated that many pathways modulated by dopamine involve IL-1β, including IL-10 Signaling, Protein Kinase A Signaling, and PPARα/RXRα Activation. We also saw decreases in several inflammatory pathways involving these same targets, including Neuroinflammation Signaling, IL-8 Signaling, iNOS Signaling, PI3K/AKT Signaling, HIF1α Signaling, Role of PKR in Interferon Induction and Antiviral Response, Chemokine Signaling, NF-κB Activation by Viruses, and IL-1 Signaling. Mapping these changes in gene expression onto functions revealed indicators of upstream regulators associated with neuronal activity and neurotransmission, microglia-enriched transcription factors and inflammatory signaling. Following up on these targets will be important to better understand mechanisms by which dopaminergic dysregulation could influence inflammatory neuropathologies. This could further guide research to better define which dopaminergic medications would mitigate excess inflammation, while sustaining protective inflammatory responses. The contrasting effects of dopamine on pro- and anti-inflammatory responses indicate that modulation of dopaminergic activity could both initiate and inhibit inflammatory cascades. Small changes in dopamine concentrations may shift activity in either direction, depending on the receptor expression on the cell type exposed to dopamine, converting a protective effect into something pathologic. The specific, and often opposing, myeloid signatures identified in the hyperdopaminergic environment modeled here suggest that increased exposure to dopamine could modulate IL-1β-mediated inflammatory signaling in human myeloid cells, suggesting an explanation for the variability in inflammatory outcomes in the context of substance use and dopaminergic medication regimens (Kohno, Link et al. 2019, Ermakov, Melamud et al. 2023, Patlola, Donohoe et al. 2023).

An example of a neuroinflammatory disease in which there is often dopaminergic dysregulation is HIV, as the prevalence of substance use disorders in people living with HIV is very high, particularly in older adults (Hartzler, Dombrowski et al. 2017, Shiau, Arpadi et al. 2017, Deren, Cortes et al. 2019). Further, HIV infection itself has been shown to dysregulate the dopaminergic system (Larsson, Hagberg et al. 1991, Aylward, Henderer et al. 1993, Berger, Kumar et al. 1994, Lopez, Smith et al. 1999, Wang, Chang et al. 2004, Kumar, Fernandez et al. 2009, Scheller, Arendt et al. 2010). These data show that the effects of dopamine on IL-1β persist and may be exacerbated in the presence of HIV infection in hMDM, C20 microglia, and iPSC-derived microglia. Notably, these data showed that HIV infection did not increase IL-1β production relative to uninfected cells, instead mostly producing non-significant decreases in baseline IL-1β levels (not shown). This comports with our prior observations regarding cytokine production in hMDM from ART-treated, HIV-infected individuals (Nolan, Muir et al. 2018), as well as other studies (Molina, Schindler et al. 1990, Brown, Kohler et al. 2008, Rozmyslowicz, Murphy et al. 2010), although are also contrasting reports indicating that HIV infection increases macrophage inflammation (Folks, Justement et al. 1987, Pontillo, Silva et al. 2012, Guo, Gao et al. 2014, Feria, Taborda et al. 2018). These differences could be due to how long after infection cytokine production was assessed, as the timing of the immune stimulus – in this case HIV infection - is very important for IL-1β production in myeloid cells (Sanz and Di Virgilio 2000). For example, peak IL-1β production in another study using HIV-infected, iPSC-derived human microglia occurred at 16 days post infection (Boreland, Stillitano et al. 2024), while in primary human microglia, HIV infection increased IL-1β release within 24 hours post-infection, which preceded peak production and release of virus (7-10 days post-infection) (Walsh, Reinke et al. 2014). This suggests that future examination of IL-1β production over time could uncover more robust interactions with dopamine. Importantly, these studies did not use anti-retroviral drugs despite the prevalence of anti-retroviral therapy (ART) in the infected population, because the impact of ART on pro-inflammatory cytokine production, such as IL-1β, is not well defined and inconsistent (Krebs, Slike et al. 2016, Okay, Koc et al. 2020, Masyuko, Page et al. 2021, Wilkinson, Schneider-Luftman et al. 2021). Future studies will need to better define the impact of ART on dopaminergic inflammation in HIV-infected individuals, but our prior data indicate that ART does not block the effects of dopamine on inflammation (Nolan, Muir et al. 2018), and the literature suggests that ART does not return inflammation to baseline levels (Regidor, Detels et al. 2011, Funderburg 2014, Sereti, Krebs et al. 2017).

Beyond the effects on IL-1β, the interactive effects of dopamine and HIV on inflammasome activity were also examined in hMDM and C20 microglia. In both HIV-infected macrophages and C20 microglia, dopamine significantly increased gene and/or protein expression of NLRP3, correlating well with our data in uninfected myeloid cells in this and our previous studies, as well as other reports showing HIV-infection is associated with elevated NLRP3 activity (Chivero, Guo et al. 2017, Bandera, Masetti et al. 2018, Feria, Taborda et al. 2018, Nolan, Reeb et al. 2020). However, dopamine had different effects on other inflammasomes in these two cell types, increasing NLRC4 and AIM2 in macrophages, and decreasing NLRP1, with no effect on other inflammasomes, in C20 cells. The different effects of dopamine on discrete types of inflammasomes could be due to the fact that dopamine may not act as the same type of stimuli for all inflammasomes, as certain stimuli are required to activate each inflammasome (Zheng, Liwinski et al. 2020). Some of these differences could also be cell type-specific, as there are differences in inflammasome expression and inflammasome-induced IL-1β secretion among myeloid cells. For example, while caspase 1, 4, and 5 activation is pivotal for inflammasome-induced IL-1β secretion by hematopoietic macrophages, microglial secretion of IL-1β is only partially dependent on these inflammatory caspases (Burm, Zuiderwijk-Sick et al. 2015). The same study found that microglia expressed significantly lower levels of AIM2 and caspase 1, 3, 4, and 5-encoding mRNA transcripts compared to BMDMs at baseline (Burm, Zuiderwijk-Sick et al. 2015). Changes in inflammasome activation and IL-1β processing can also be mediated by a number of other mechanisms including P2X7 receptor activity (Oliveira-Giacomelli, Petiz et al. 2021), activation of caspase-3 or -8 inflammasomes (Burguillos, Deierborg et al. 2011, Gringhuis, Kaptein et al. 2012, Antonopoulos, Russo et al. 2015, Burguillos, Svensson et al. 2015), and several inflammasome-independent mechanisms, such as matrix metalloproteinases (Schönbeck, Mach et al. 1998), cathepsins (Orlowski, Colbert et al. 2015, Chevriaux, Pilot et al. 2020), and serine proteases (Karmakar, Sun et al. 2012, Schreiber, Pham et al. 2012, Rodriguez, Ducker et al. 2019).

There are also multiple types of inflammasomes that can influence production of other IL-1 cytokines, such as IL-18. Surprisingly, in our HIV-infected C20 microglia, the effects of dopamine on IL-18 were dissimilar to IL-1β in that there was a trend towards a decrease in IL-18 expression. Part of this may be due to the fact that IL-18 has been shown to be constitutively expressed in myeloid cells (Ahmad, Sindhu et al. 2002, Dinarello, Novick et al. 2013). In contrast, IL-1β is not constitutively expressed under homeostasis and requires induction with TLR ligands or other stimuli. Additionally, IL-18 is considered an inflammasome-regulated cytokine but reports have shown it is only marginally influenced by LPS and ATP at both mRNA and secreted protein levels, suggesting that its modulation is not similar to IL-1β, and additional pathways may regulate its levels (Awad, Assrawi et al. 2017). Indeed, there are several reports for the role of NLRP1 in IL-18 induction (Nour, Yeung et al. 2009, Murphy, Kraakman et al. 2016), which may explain their decreased expression. Further, although inflammasome signaling in the CNS is mainly attributed to microglia, expression of inflammasome components has also been reported in other cell types of the CNS, including perivascular CNS macrophages and astrocytes (Kawana, Yamamoto et al. 2013), so important additional cell type-specific differences in inflammasome-mediated responses could contribute to disease pathology (Voet, Srinivasan et al. 2019).

These data highlight the potential significance of developing a better understanding of dopaminergic immunomodulation in myeloid cells. However, there are caveats that must be considered when moving this research forward. First, these studies focused on dopamine receptor-mediated effects, as recent studies have also shown that other dopaminergic proteins, such as the dopamine transporter and tyrosine hydroxylase, can also regulate myeloid inflammation (Mackie, Gopinath et al. 2022) and alter the peripheral immune profile in dopaminergic diseases such as Parkinson’s disease (Gopinath, Mackie et al. 2022). Another consideration for future studies are the cell models being used, as several limitations for these studies are the limited number of human myeloid cell lines, the lack of access to primary human tissue macrophages – such as microglia - and the well-known variability in primary human monocyte derived macrophages (Bol, van Remmerden et al. 2009, Jobe, Kim et al. 2017). To address the heterogeneity in the human population, we used hMDM from many donors (n = 42) relative to many *in vitro* studies, but the number was still small compared to epidemiological studies examining trends across a human population. For example, for the dopamine receptor combination analyses in **Table 1**, there were several trends that might have shown significance if the experiments were powered to assess these questions. It is also important to note that the PBMC / monocyte / hMDM populations described in **Figure 2** and **Supplementary Figure 3** were not matched for every donor, as matched samples were not always available. However, analysis of only the matched PBMC / monocyte / hMDM donor sets (n = 18 - 19) showed the same effects, strengthening confidence in these data and strongly indicating that dopamine receptor expression, at least in human myeloid cells, varies with cell type and maturation.

Because acquisition of primary human microglia is logistically difficult, and because cell lines provide a more consistent experimental platform than primary macrophages, established C06 and C20 immortalized human microglial cells (Garcia-Mesa, Jay et al. 2017, Alvarez-Carbonell, Ye et al. 2019, Ye, Alvarez-Carbonell et al. 2022, Rheinberger, Costa et al. 2023) were used for many experiments. These cells showed a clear, replicable dopamine response, but they also show limitations, as they replicate rapidly and do not recapitulate the viral dynamics in primary myeloid cells. Further, we were only able to detect IL-1β transcripts and intracellular IL-1β production, not IL-1β secretion, in these cells. We also could not detect caspase-1, so it is possible other caspases were involved in this response. To address these concerns, studies were performed across three distinct human myeloid model systems, all of which showed the same effects of dopamine. Much of the research on dopaminergic immunology has been performed in rodent models, making the use of multiple human myeloid models one of the major strengths in this study. Further studies in human systems, as well as correlation of the extant rodent research with human data is critical to address issues of translatability arising from species-specific differences in rodent and human immune responses and inflammasome activity (Mestas and Hughes 2004, Ariffin and Sweet 2013, Shay, Jojic et al. 2013, Schaale, Peters et al. 2016, Egan, Zhang et al. 2023), which may, in part, contribute to the difficulty translating therapeutics from preclinical models (Copeland, Warren et al. 2005, Lamkanfi and Dixit 2012, Gosselin, Skola et al. 2017). Similar issues could be caused by differences in dopamine receptor ratios, which also differ across species (Mandeville, Sander et al. 2013), as well cell-type specific differences in dopamine signaling activity (Nickoloff-Bybel, Mackie et al. 2019). Adding further complexity is the fact that higher concentrations of dopamine can also activate α- and β-adrenergic receptors (Lei 2014, Parrado, Salaverry et al. 2017, Sánchez-Soto, Casadó-Anguera et al. 2018). These receptors are also expressed in human and rodent immune cells, with relative ratios of adrenergic receptors to dopamine receptors that differ across, species, cell type and brain region (Goldman-Rakic, Lidow et al. 1992, Sánchez-Soto, Casadó-Anguera et al. 2018, Aslanoglou, Bertera et al. 2021). As the data presented here suggest that IL-1β regulation may relate to differential expression and regulation of dopamine receptors, it is critical to consider these factors to define the inflammatory impact of dopamine more comprehensively.

The inflammatory impact of dopamine could influence a wide array of pathological conditions as it dysregulates inflammation in myeloid cells across regions. Many peripheral tissues with high levels of dopamine harbor one or more types of hMDM populations (Mass, Nimmerjahn et al. 2023), and it is not clear how the effects of dopamine differ within these populations. However, these data show that there can be substantial differences between myeloid cell types. Thus our ongoing research is interrogating the effects of dopamine on inflammasome activity and immune responses in tissue-specific myeloid cell types. Many of these effects could be exacerbated in the presence of diseases, substances of misuse and therapeutics that dysregulate the dopaminergic system (Gaskill and Khoshbouei 2022). For example, a number of neuropsychiatric diseases associated with altered dopaminergic neurotransmission are also associated with inflammation, and a number of neurotransmitter-altering therapeutics used to treat these diseases also impact inflammation (Kubera, Kenis et al. 2004, Buttarelli, Fanciulli et al. 2011, Hannestad, DellaGioia et al. 2011, Hiles, Baker et al. 2012, Nazimek, Strobel et al. 2017, Kohler, Freitas et al. 2018, Wiedlocha, Marcinowicz et al. 2018, Matt and Gaskill 2019, Matt 2021, Channer, Matt et al. 2022). It is possible that the changes in the dopaminergic system resulting from either the disease or the treatment could create a vicious cycle, altering myeloid activity to exacerbate disease, reducing or counteracting the effectiveness of treatment, and promoting further dysregulation that drives the cycle forward. A similar effect could be produced in neuroHIV, where the aging, HIV-infected population is frequently prescribed dopamine-modulating therapeutics for age-associated comorbidities, such as Alzheimer’s, Parkinson’s, diabetes, and some cancers (Matt and Gaskill 2019).

With a better understanding of the inflammatory processes driven by dopamine, such as the interaction between dopamine receptor-mediated signaling and IL-1β, a combination of targeting IL-1β alongside dopaminergic machinery could be a promising avenue to prevent or limit widespread chronic immune activation associated with numerous pathologies. Overall, these data demonstrate the remarkable complexity of dopaminergic immunomodulation and indicate that more research is needed to better understand how these effects are regulated, how they differ throughout the body, and how best to use this knowledge to repurpose or develop new medication regimens in the context of HIV and other brain diseases associated with chronic inflammation.

## Conflict of Interest

The authors declare that the research was conducted in the absence of any commercial or financial relationships that could be construed as a potential conflict of interest.

## Author Contributions

SMM, RN, and PJG contributed to the design and conception of the study. SMM, RN, BL, MN, JS, HF, and PJG designed and analyzed the experiments, and SMM, RN, SM, YA, BC, OT, MD, JC, TL, KM, KR, ENB, BL, and JS performed the experiments. SMM and PJG helped supervise the project. SMM and PJG performed the statistical analyses and wrote the manuscript. All authors contributed to manuscript revision, read, and approved the final submitted version, and PJG was responsible for the final approval of the submitted version.

## Funding

This work was supported by grants from the National Institutes of Drug Abuse, DA039005, DA057337, DA058051 and DA049227 (PJG), the W.W. Smith Charitable Trust Foundation Grant A2003 (PJG), MH132466 (SMM), the Brody Family Medical Trust Fund (SMM), the Cotswold Foundation Fellowship (SMM), T32-MH079785 (supporting RAN, EAN, BC) and the Department of Pharmacology and Physiology at Drexel University College of Medicine.

## Acknowledgments

We would like to state our tremendous appreciation for all the individuals who donated the biological materials used in these studies. We would also like to thank all the members of the Gaskill lab as well as Dr. Vasiliki Pappa, Katelyn Reeb, and John Montilla for their contributions to this work.

## Notes

### Competing Interest Statement

The authors have declared no competing interest.

