## Supplementary material for "Dopamine-driven Increase in IL-1β in Myeloid Cells is Mediated by Differential Dopamine Receptor Expression and Exacerbated by HIV": Supp Figures and Supp Table 1

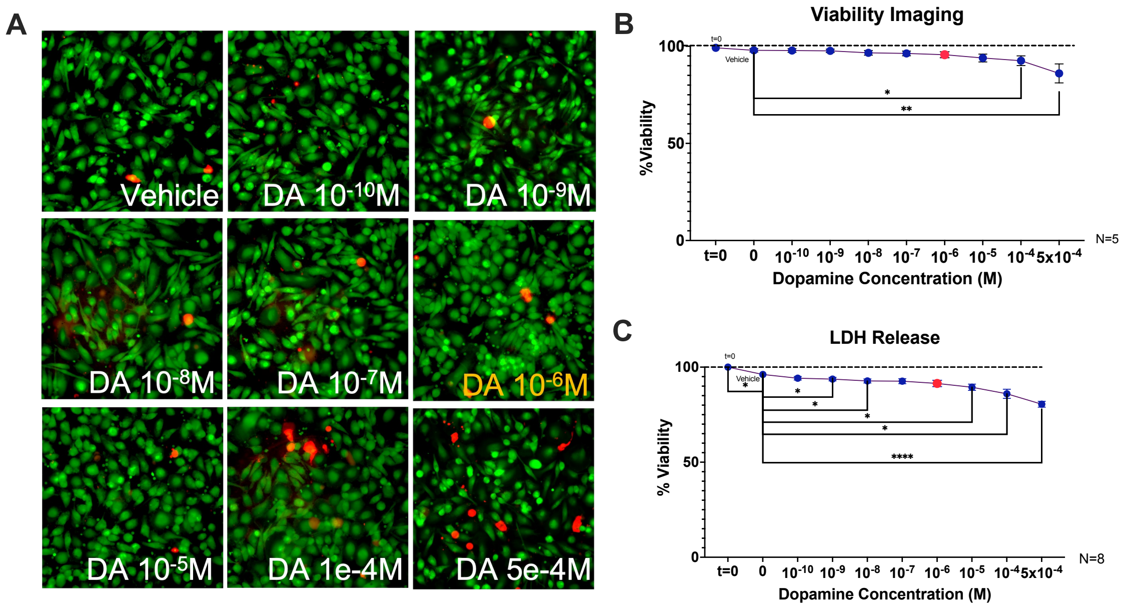


**Supplementary Figure 1. 1x10$^{-6}$ M Dopamine does not affect primary human macrophage viability. (A)** Representative high content images (20X objective) of primary human monocyte derived macrophages treated for 24 hours with vehicle or dopamine conditioned media (1x10$^{-10}$ to 5x10$^{-4}$M) and stained, using LIVE (green) DEAD (red) cell imaging kit. **(B)** Quantification of high content imaging of viability assay (N=5). **(C)** LDH release from supernatants from 8 independent experiments of 24hr incubation with and without Dopamine (1x10$^{-10}$ to 5x10$^{-4}$M). Significance was determined using Friedman, one-way rm ANOVA and paired t-tests, *p < 0.05, **p < 0.01, and ****p < 0.0001. The concentration used in all subsequent studies (10$^{-6}$M ) is denoted in orange in **A** and red in **B-C**.

### Supplementary Table 1: Parameters for high content analysis of immunocytochemistry

| Condition | Value |
|---|---|
| Ch1 (LIVE): Smoothing: Uniform | NA |
| Ch1 (LIVE): Thresholding (Fixed) | 500 |
| Ch1 (LIVE): Segmentation (Intensity) | 500 |
| Ch1 (LIVE): Object Cleanup | Y |
| Object.BorderObject.Ch1 | Y |
| Object.Ch1.Area.Ch1 | 854.84-7746.16 |
| ObjectCh1.ShapeP2A.Ch1 | NA |
| ObjectCH1.ShapeLWR.Ch1 | NA |
| Object.AvgIntensity.Ch1 | NA |
| Object.TotalIntensity.Ch1 | NA |
| Object.VarIntensity.Ch1 | 7.52-65535 |
| Ch2 (DEAD): Thresholding (Fixed) | 2000 |
| Ch2 (DEAD): Segmentation (Intensity) | 28 |
| Object.BorderObject.Ch2 | Y |
| Object.Ch2.Area.Ch2 | 265.28-6653.4 |
| Object.AvgIntensity.Ch2 | NA |
| Object.TotalIntensity.Ch2 | NA |
| Object.VarIntensity.Ch2 | NA |
| ObjectCh2.ShapeP2A.Ch2 | NA |
| ObjectCH2.ShapeLWR.Ch2 | NA |
| ROIA.Mask.Ch1 | Ch1 (Green Live) |
| ROIA.Target1 | Ch2 (Red Dead) |

**Legend:** Y = yes or selected; NA = Not applicable

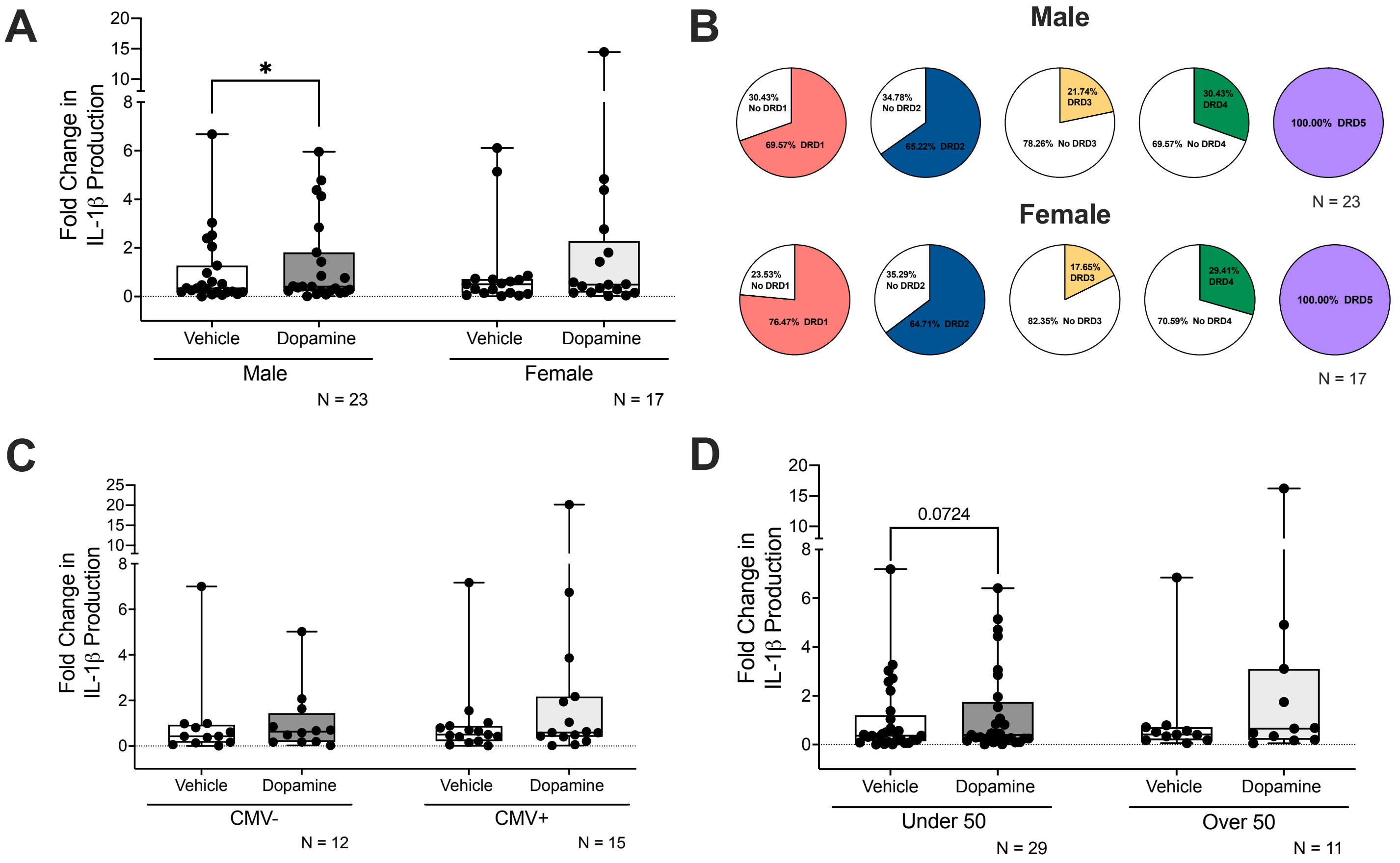


**Supplementary Figure 2. Dopamine-mediated changes in IL-1β production associated with donor demographic data.** Analyses were performed to look at dopamine-mediated changes in IL-1β in association with demographic data. **(A)** Males had a significant dopamine-mediated increase in IL-1β while females did not. However, this cannot be explained by differences in distribution of dopamine receptor subtypes between males and females, as there were no statistically significant alterations in expression of any of the dopamine receptors **(B)**. **(C)** Neither CMV+ nor CMV- individuals had dopamine-mediated increases in IL-1β. **(D)** There was a trend in that individuals under 50 years old had a dopamine-mediated increase in IL-1β. Significance was determined using Wilcoxon tests, *$p < 0.05$.

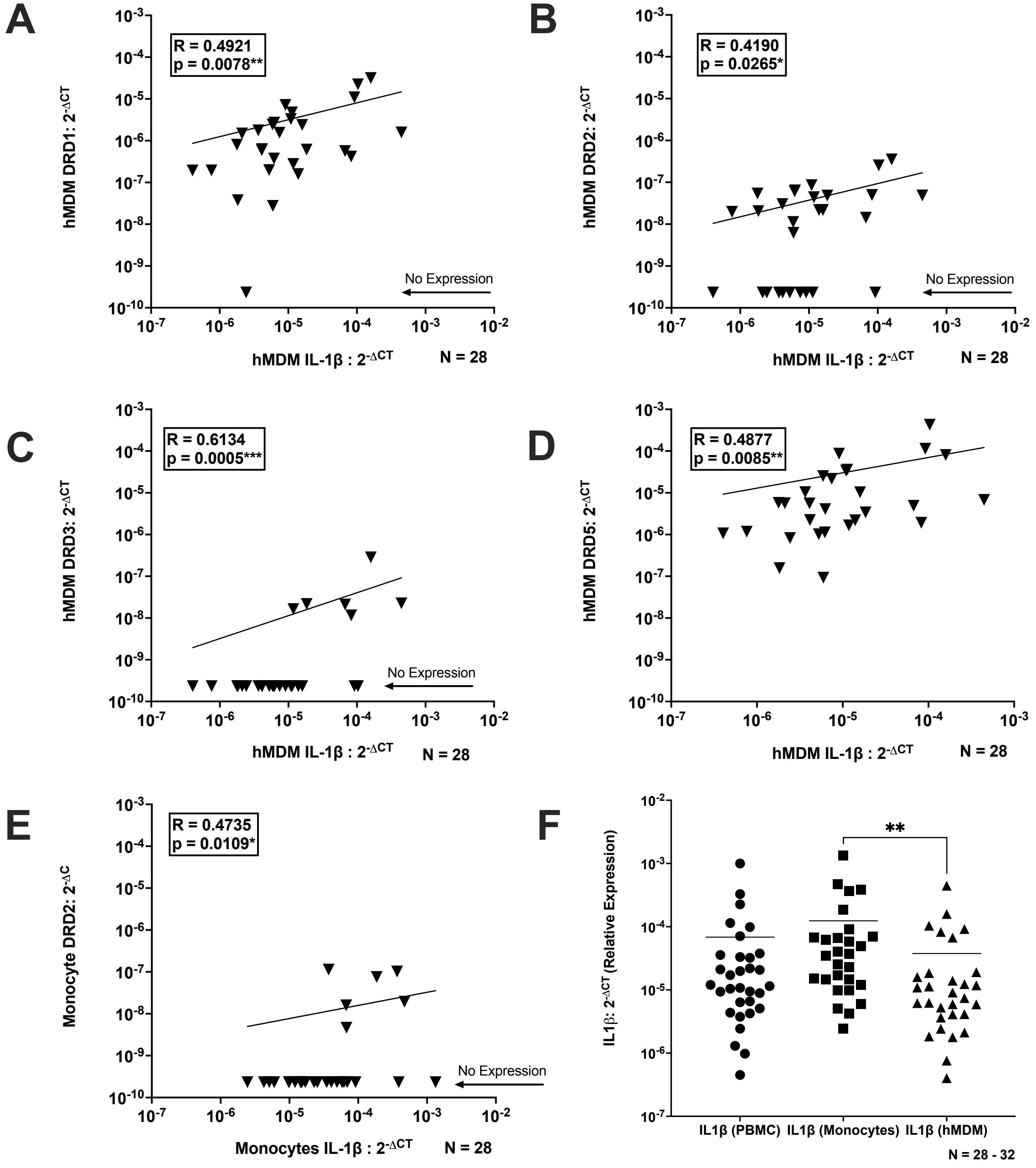


**Supplementary Figure 3: IL-1β expression levels in primary human myeloid cells differentially correlate with dopamine receptors.** Data were analyzed for correlations between IL-1β and each dopamine receptor between primary human monocyte-derived macrophages (hMDMs), monocytes, and peripheral blood mononuclear cells (PBMCs). These analyses showed significant positive correlations between IL-1β and **(A)** DRD1, **(B)** DRD2, **(C)** DRD3, and **(D)** DRD5 in hMDMs. There was only a significant positive correlation between IL-1β and DRD2 in monocytes **(E)**, and no correlations between IL-1β and dopamine receptors in PBMCs. **(F)** Baseline IL-1β mRNA was compared between the three cell types (18 of which were paired for all three cell types) and monocytes had significantly higher baseline IL-1β compared to hMDMs. Significance was determined using Spearman correlations, Kruskal-Wallis tests, and post-hocs with Dunn's multiple comparisons,*$p < 0.05$, **$p < 0.01$, and ***$p < 0.001$.

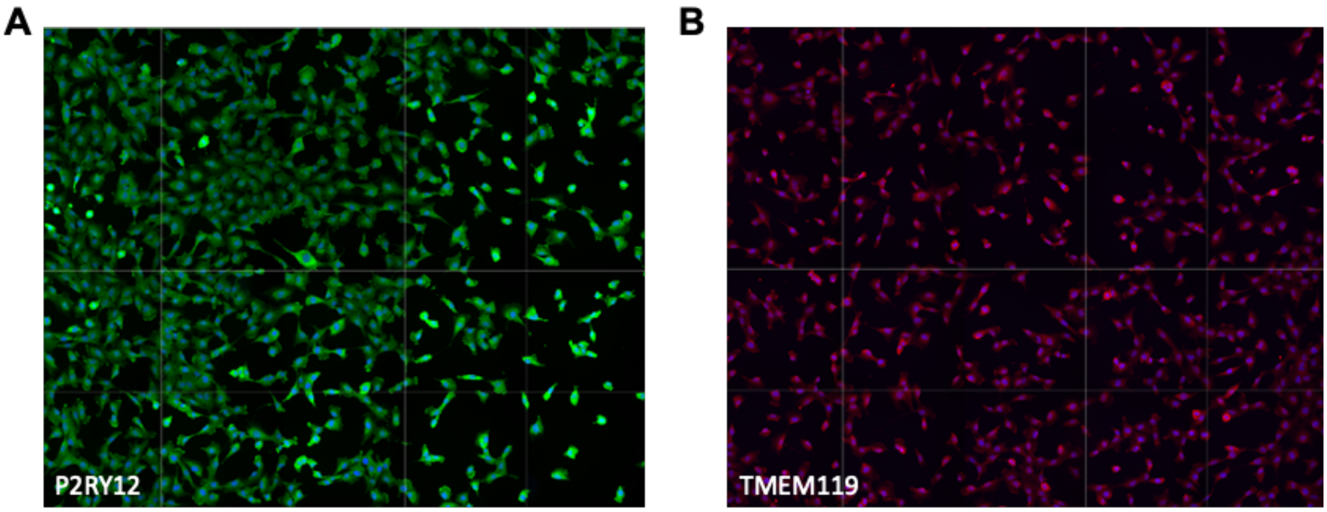


**Supplementary Figure 4: Microglial marker expression in C20 cells.** C20 microglia were plated at 5,000 cells per well in a 96-well plate and stained with DAPI and P2RY12 (1:100) or TMEM119 (1:50). Representative stitched CX7 20X objective images of C20 microglia stained with **(A)** P2RY12 or **(B)** TMEM119.

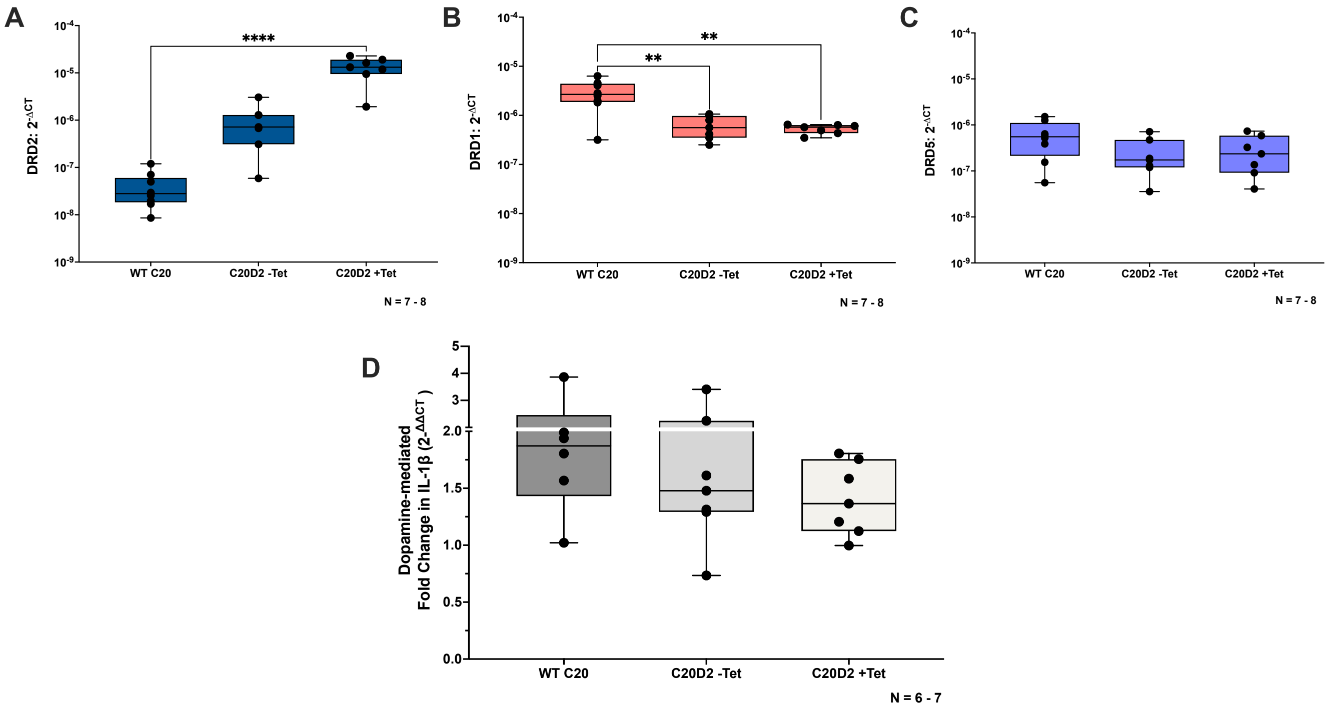


**Supplementary Figure 5: Overexpression of DRD2 in microglia dampens dopamine-induced increase in IL-1β.** In the absence or presence of tetracycline for 48 hours, C20D2 microglia were examined for dopamine receptors measured by qPCR and compared to WT. **(A)** C20D2 microglia +/- Tet expressed significantly more DRD2 **(B)** and less DRD1 than WT C20 microglia, but **(C)** there was no change in DRD5 expression. **(D)** C20D2 microglia were treated with dopamine ($10^{-6}$ M) for 3 hours and examined for IL-1β mRNA. Compared to WT C20 microglia, C20D2-Tet had a decreased dopamine-mediated fold change in IL-1β, and this was further decreased with C20D2+Tet. Significance was determined by a one-way ANOVA and post-hoc with Tukey's multiple comparisons, $^{**}p < 0.01$, $^{****}p < 0.0001$.

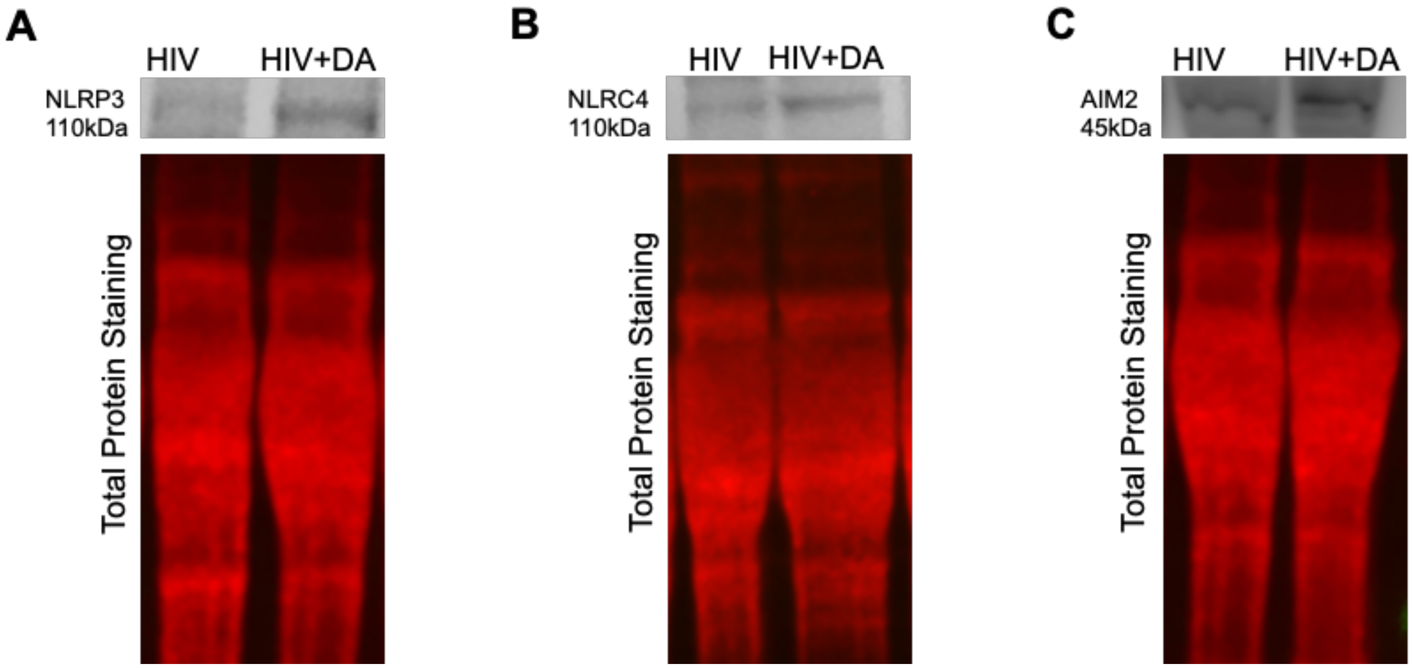


**Supplementary Figure 6. Representative Inflammasome Western Blots in hMDM.** Western blots of total protein stain (TPS) and **(A)** NLRP3, **(B)** NLRC4, and **(C)** AIM2 in hMDM.

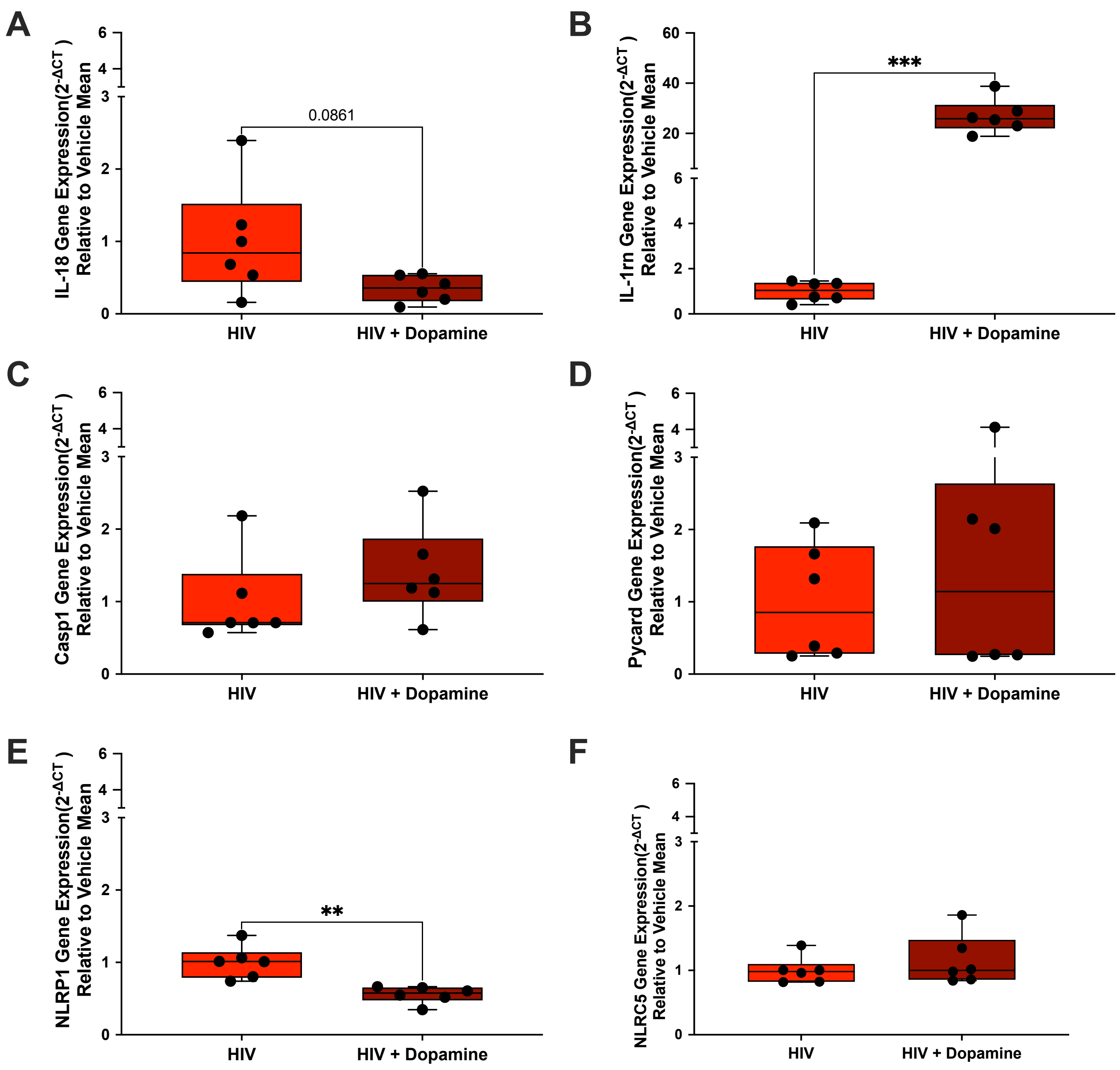


**Supplementary Figure 7: Dopamine modulates expression of IL-1-family and inflammasome components.** C20 microglia were infected with 2.5 ng/ml $HIV_{ADA}$ for 48 hours, then treated with dopamine ($10^{-6}$ M) for 3 hours and examined for **(A)** IL-18, **(B)** IL-1rn, **(C)** Casp1, **(D)** Pycard, **(E)** NLRP1, and **(F)** NLRC5 mRNA expression by qPCR. There was a trend in decreased IL-18 expression, and there was a significant increase in IL-1rn and decrease in NLRP1 with HIV + Dopamine relative to HIV alone. Significance was determined by paired t-tests and Wilcoxon tests, $^{**}p < 0.01$ and $^{***}p < 0.001$.

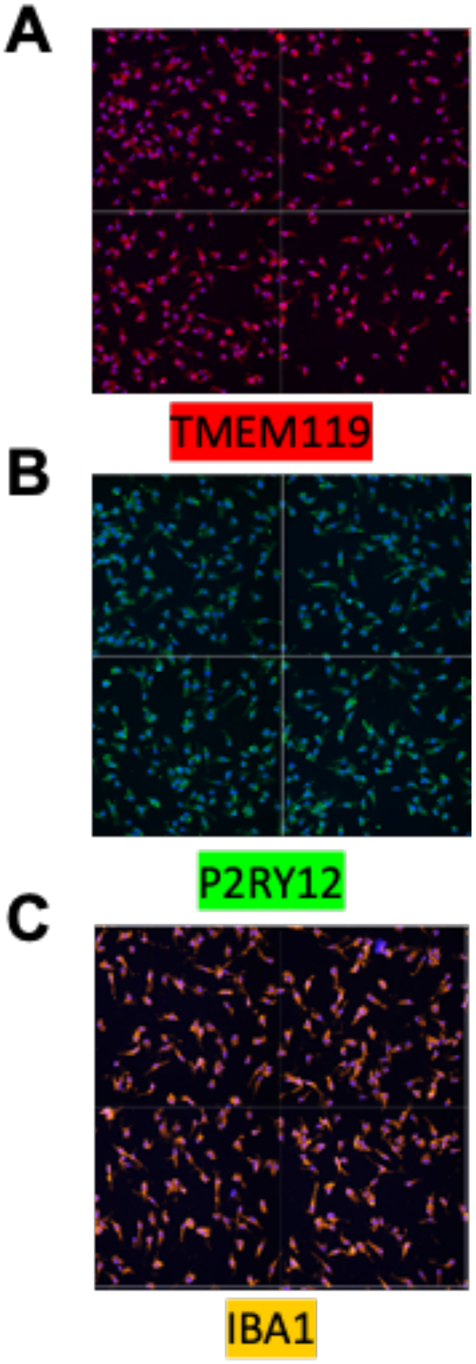


**Supplementary Figure 8: Microglial Marker Expression in iPSC-derived Microglia.** iPSC-derived microglia were plated at 5,000 cells per well in a 96-well plate and stained with DAPI and TMEM119 (1:50), P2RY12 (1:100) or IBA1 (10ug/mL). Representative stitched CX7 images of C20 microglia stained with **(A)** TMEM119, **(B)** P2RY12, or **(C)** IBA1.
